# Neuronal Activity-Dependent Gene Dysregulation in *C9orf72* i^3^Neuronal Models of ALS/FTD Pathogenesis

**DOI:** 10.1101/2025.01.27.632228

**Authors:** Layla T. Ghaffari, Emily Welebob, Ashley Boehringer, Kelly Cyliax, Piera Pasinelli, Davide Trotti, Aaron R. Haeusler

## Abstract

The GGGGCC nucleotide repeat expansion (NRE) mutation in the *C9orf72* (C9) gene is the most common cause of ALS and FTD. Neuronal activity plays an essential role in shaping biological processes within both healthy and neurodegenerative disease scenarios. Here, we show that at baseline conditions, C9-NRE iPSC-cortical neurons display aberrations in several pathways, including synaptic signaling and transcriptional machinery, potentially priming diseased neurons for an altered response to neuronal stimulation. Indeed, exposure to two pathophysiologically relevant stimulation modes, prolonged membrane depolarization, or a blockade of K^+^ channels, followed by RNA sequencing, induces a temporally divergent activity-dependent transcriptome of C9-NRE cortical neurons compared to healthy controls. This study provides new insights into how neuronal activity influences the ALS/FTD-associated transcriptome, offering a dataset that enables further exploration of pathways necessary for conferring neuronal resilience or degeneration.

## Introduction

The (GGGGCC)_n_ nucleotide repeat expansion (NRE) mutation located in the *C9orf72* gene is the most common genetic cause of Amyotrophic lateral sclerosis (ALS) and frontotemporal dementia (FTD), two devastating and progressive neurodegenerative diseases. ALS affects both the upper and lower motor neurons of the corticospinal tract, while FTD primarily affects cortical neurons of the frontal and temporal lobes. The discovery of the NRE in *C9orf72* has elucidated that ALS and FTD are part of an overlapping disease spectrum (Ling et al., 2013). The presence of the NRE is proposed to lead to neurodegeneration in three ways. Firstly, the NRE in intron 1 of *C9orf72* impairs transcription of the NRE-affected allele, and thus, there is a reduction in C9ORF72 protein. The full impact of the loss of C9ORF72 remains obscured, however, in induced pluripotent stem cell (iPSC) motor neuron models, a loss of C9ORF72 renders motor neurons susceptible to death by secondary insults such as glutamate excitotoxicity or reduction in neurotrophic factors (Shi et al., 2018). Secondly, the NRE is transcribed into repeat-containing RNA, which forms RNA foci and sequesters essential RNA-binding proteins which impairs RNA metabolism (DeJesus-Hernandez et al., 2011; Donnelly et al., 2013; Lee et al., 2013; Haeusler et al., 2014; Conlon et al., 2016). Thirdly, repeat-containing RNA can be aberrantly translated in a non-canonical manner called repeat-associated non-AUG translation (RAN) into 5 distinct dipeptide repeat protein (DPRs) species. DPRs have been proposed to exert neurotoxic effects, dependent upon DPR species, localization, and length (DeJesus-Hernandez et al., 2011; Renton et al., 2011; Mackenzie et al., 2013; Freibaum and Taylor, 2017). The NRE in the *C9orf72* gene is associated with synaptic dysfunction, however, the precise relationship remains unclear (Ghaffari et al., 2022). In a healthy context, the activity-dependent gene transcription pathway is critical in synaptic remodeling, neuronal health, and responding to cellular stimuli (Flavell and Greenberg, 2008; Kim et al., 2010; Guo et al., 2011; Benito and Barco, 2015; Su et al., 2017; Yap and Greenberg, 2018). Outside of a healthy context, it is known that deficits in the activity-dependent transcriptome can result in neurodevelopmental deficits, intellectual disability, and diseases affecting cognition (Ebert and Greenberg, 2013; Yap and Greenberg, 2018). Although these neurological diseases typically manifest earlier in life, in contrast with neurodegenerative disease, the evidence for neuronal-specific mechanisms of dysfunction suggests their potential importance in age-related disorders. It has been established that altered neuronal activity, in the form of cortical hyperexcitability, is present in ALS patients early in the disease manifestation (Vucic et al., 2008; Kiernan, 2009; Bae et al., 2013; Wainger and Cudkowicz, 2015; Schanz et al., 2016). This early altered neuronal activity has been proposed to be an initiating event, contributing to the “dying-forward” hypothesis of ALS, with disease arising in the corticomotor neurons and disease pathogenesis descending the corticospinal tract (Kiernan et al., 2011). The link between neuronal activity and disease-relevant events such as nuclear loss of TDP-43 (Weskamp et al., 2020) and elevation of dipeptide repeat proteins (DPRs) (Westergard et al., 2019; Catanese et al., 2021) is also unclear.

We recently reported that neuronal depolarization modulates C9orf72 transcript variant levels in healthy human induced pluripotent stem cell cortical-like neurons (i^3^Neurons), and the same transcripts are differentially regulated in C9-NRE carrier-derived iPSC i^3^Neurons (Ghaffari et al., 2023). These intriguing results led us to question if the widespread transcriptomic landscape of C9-NRE i^3^Neurons may have an altered response following neuronal depolarization or increased excitation. Here, we show that C9-NRE i^3^Neurons display unique temporal gene expression profiles following neuronal depolarization and increased neuronal excitation. This leads to the emergence of several overlapping and distinct pathways that could be critical in disease pathogenesis. Furthermore, some of these pathways, such as synaptic programming, validate and extend previous observations from both patient tissues and in vitro/vivo disease models, while others, such as crystallin or peroxisomal dysregulation, provide new potential insights as well as therapeutic targets for C9-NRE-linked diseases to be explored in future studies. Therefore, we have made this dataset publicly available for exploration in a Shiny application (https://haeuslerlab.shinyapps.io/ltg_shiny/).

## Results

### Dysregulation of synaptic genes in C9-NRE i^3^Neurons correlates with elevated neuronal oscillation and synchronicity

We first utilized a multielectrode array (MEA) system to evaluate potential baseline differences in neurophysiological properties for i^3^Neurons generated from healthy individuals versus C9-NRE ALS/FTD patients. i^3^Neurons are a human induced pluripotent stem cell (hiPSC) model in which hiPSC lines have a doxycycline-inducible Neurogenin-1 and/or Neurogenin-2 (NGN1/NGN2) cassette integrated into a safe-harbor locus site (AAVS1 or CLYBL) (Fernandopulle et al., 2018; Weskamp et al., 2020). Doxycycline addition leads to hiPSC differentiation into glutamatergic cortical-like neurons in a quick and reproducible manner. We used i^3^Neurons lines derived from healthy individuals and C9-ALS/FTD patients (C9-NRE hereafter). The three major categorical readouts from MEA recordings are neuronal activity, oscillation, and synchrony (**Figure 1A**). Neuronal activity serves as a validation of functional neurons, and a measure of this is weighted mean firing rate (Hz), which is the number of spikes throughout the recording, as measured by active electrodes. As previously reported, we did not observe significant differences in mean firing rates between control and C9-NRE i^3^Neurons on the day of experimentation (DIV 25) (**Figure 1B**) or at any time during the differentiation to DIV 25 (Ghaffari et al., 2023). Measuring network functionality can be done through readouts of oscillation, such as network burst frequency and number of network bursts, which reveal the organization of network activity. There is a slight but insignificant increase in the Network Burst Frequency (Hz) and Number of Network Bursts of C9-NRE i^3^Neurons at DIV 25 (**Figure 1B**). Synchrony is a measure of functional synapses and, in short, neurons are considered synchronous if spikes from one electrode led to spikes in a neighboring electrode. The Synchrony Index measurement ranges from 0-1, with 0 indicating no synchrony and 1 indicating complete synchrony. The C9-NRE i^3^Neurons show slightly increased synchronicity compared to Controls, as shown by the Synchrony Index and Area Under Normalized Cross Correlation (**Figure 1B**). These findings demonstrate that C9-NRE i^3^Neurons do not overtly show hyperexcitable phenotypes but have slightly more network activity and display slight increases in synchrony. These phenotypes, while subtle, may point to dysregulation of synaptic gene networks that could be more sensitively captured by transcriptomic analysis.

**Figure 1.**
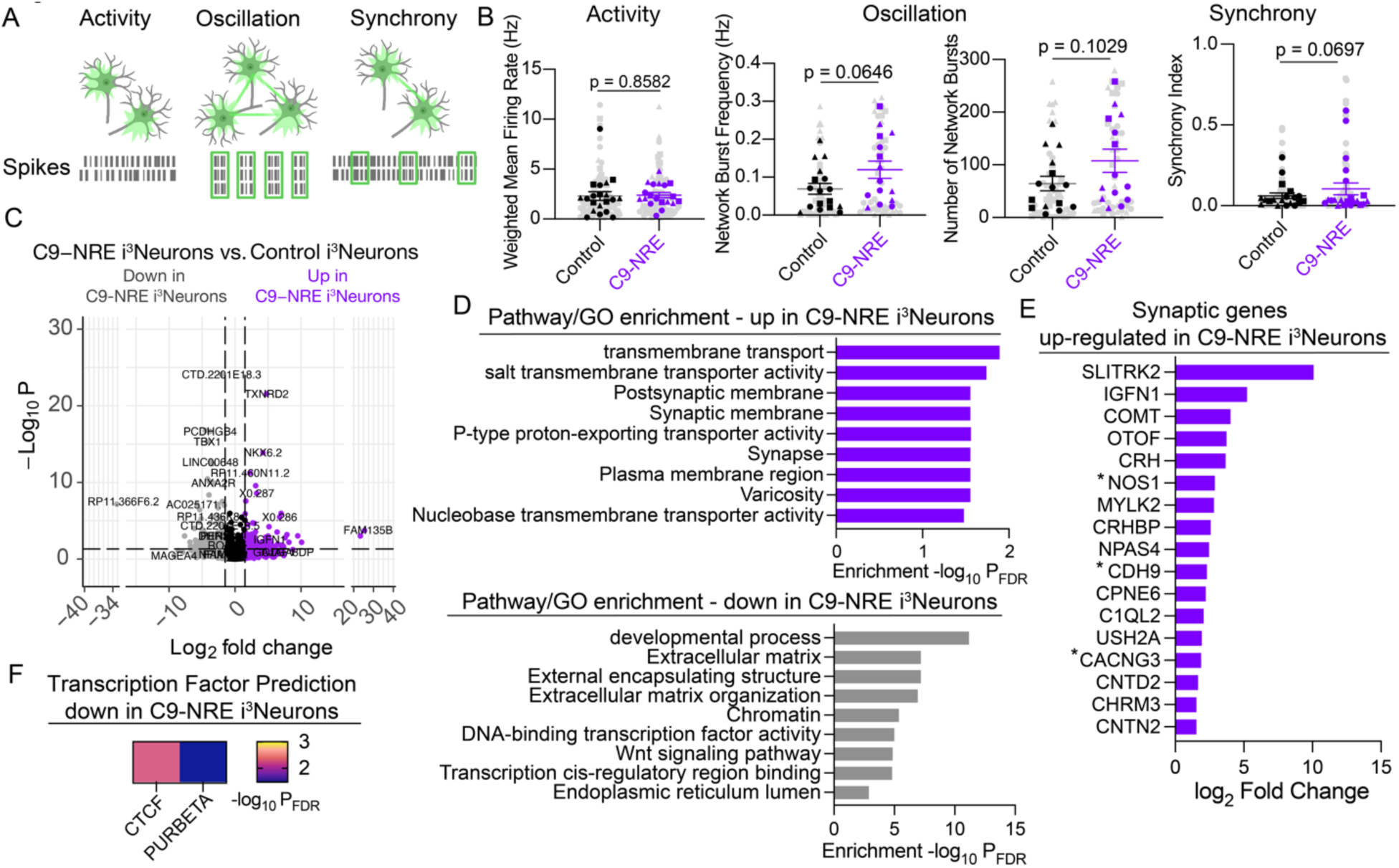
C9-NRE i^3^Neurons are slightly more synchronous and display aberrant levels of synaptic transcripts. **(A)** Schematic of the metrics from multi-electrode array recordings. Activity refers to functional readout, oscillation refers to the frequency and pattern of firing, and synchrony refers to the connection between neurons on the electrodes. **(B)** Weighted mean firing rate is a measure of activity. There are no differences in the weighted mean firing rate of Control and C9-NRE i^3^Neurons at DIV 25. Network Burst Frequency measured in Hertz (Hz) over the time of the recording shows a slight and nearly significant increase in C9-NRE i^3^Neurons. The total number of Network Bursts is trending upward in the C9-NRE i^3^Neurons. Synchrony of C9-NRE i^3^Neurons, as plotted by the Synchrony Index, shows a nearly significant increase compared to Controls. For each experiment, *n* = 3 biological replicates, *m* = 4-9 MEA plates, *o* = 4-6 wells per plate. Each biological replicate is indicated by a unique shape. Replicates on each plate were collapsed. Unpaired t-tests were performed to determine significance. Error bars represent the SEM. **(C)** Volcano plot of differentially expressed genes (DEGs) in the comparison between untreated Control i^3^Neurons and untreated C9-NRE i^3^Neurons. For each experiment *n* = 3 biological replicates, *m* = 2 technical replicates. **(D)** Enriched Gene Ontology (GO) terms and KEGG pathways for genes upregulated in C9-NRE and Controls are plotted. The log2 fold change of synaptic genes identified to be upregulated in C9-NRE i^3^Neurons is plotted. Predicted regulators of DEGs in Control i^3^Neurons are plotted. **(F)** Predicted transcription factor activity is based on differentially expressed genes downregulated in C9-NRE i^3^Neuron.

To understand the transcriptomic differences between Control and C9-NRE i^3^Neurons with altered neurophysiological readouts and performed bulk RNA sequencing on DIV 25 Control and C9-NRE i^3^Neurons. Differential gene expression comparing untreated (UT) Control i^3^Neurons to UT C9-NRE i^3^Neurons (**Figure 1C**) revealed an upregulation of genes enriched for synaptic gene ontology (GO) terms in C9-NRE i^3^Neurons (**Figure 1D**). Among these genes is SLITRK2, a post-synaptic transmembrane protein, (**Figure 1E**), previously reported to be upregulated in C9-ALS motor neurons and patient prefrontal cortex tissue and shown to be a genetic modifier of C9-associated FTD (Sareen et al., 2013; Satoh et al., 2014; Barbier et al., 2021). Other synaptic genes were upregulated in C9-NRE i^3^Neurons, including IGFN1, COMT, and CACNG3 (**Figure 1E**). In line with altered excitability properties, CACNG3, a voltage-gated calcium channel subunit, is linked with epilepsy (Everett et al., 2007), but also functions to decrease AMPA receptor deactivation (Kato et al., 2010). Strikingly, CACNG3 protein was also upregulated in the prefrontal cortex of C9-NRE carriers when compared to C9-NRE negative ALS patients, indicating a potentially C9-NRE specific alteration (Laszlo et al., 2022). Other genes upregulated in C9-NRE i^3^Neurons are enriched for the “transmembrane transport” GO term, which encompasses some members of the solute carrier family and ATP-ases including SLC26A9, SLC28A2, SLC5A11, SLC30A8, SLC17A9, SLC23A1, ATP12A, and ATP4A. The immediate early gene and activity-dependent transcription factor NPAS4 is upregulated at baseline in the C9-NRE i^3^Neurons. Upon examining the target genes of NPAS4 (Pollina et al., 2023), we found that several synaptic genes that are upregulated in C9-NRE i^3^Neurons are indeed targets of NPAS4, including NOS1, CDH9, and CACNG3 (denoted by *) (**Figure 1E**). Additionally, the activity-dependent transcription factor MEF2C is significantly downregulated in C9-NRE i^3^Neurons but is also an NPAS4 target gene. These findings indicate the potential for widespread NPAS4 dysregulation. Genes upregulated in Control i^3^Neurons (downregulated in C9-NRE i^3^Neurons) are enriched for transcriptional machinery and extracellular matrix (ECM) terms. A downregulation of ECM genes has also been observed in C9-NRE iPSC motor neurons (Satoh et al., 2014). Furthermore, genes upregulated in Control i^3^Neurons are predicted to be regulated by CTCF1, a chromatin architecture regulator crucial for remote memory and cortical synaptic plasticity (Kim et al., 2018). In addition to MEF2C, the transcriptional machinery-related genes that are dysregulated include players of the FOX (FOXD1, FOXF2) and HOX (HOXB2, HOXB3) family of transcription factors. Collectively, these findings show subtle alterations in synchrony and oscillation as well as disrupted synaptic transcriptomic signatures that are present in C9-NRE i^3^Neurons.

### Silenced neurons maintain dysfunctional synaptic signatures

We next examined if silencing neuronal activity may reveal inherent or responsive synaptic dysfunction signatures. To determine persistent dysregulation of synaptic genes in the absence of neuronal activity, we silenced endogenous synaptic activity with tetrodotoxin (TTX) for 24 hours, eliminating any differences in neuronal activity, oscillation, and synchrony metrics. We then compared the transcriptomes of TTX-silenced control neurons to TTX-silenced C9-NRE i^3^Neurons (**Figure 2A**). Pathway and gene ontology (GO) enrichments were strongly enriched for synaptic terms in this comparison (**Figure 2B**). This comparison revealed the persistent upregulation of synaptic genes CACNG3, NOS1, and CDH9 independent of synaptic activity. Another gene, NPAS4, known to be upregulated by neuronal activity, and is no longer upregulated in C9-NRE i^3^Neurons following TTX silencing, as expected. These results indicate that a subset of synaptic genes are differentially expressed independent of silencing induced by TTX. A further comparison of TTX-silenced Control and C9-NRE i^3^Neurons revealed additional dysregulated synaptic genes, including upregulation of GRM3, ACTC1, IL1RAP1, MCTP2, and REM2 in TTX-silenced C9-NRE i^3^Neurons (**Figure 2C,E**). REM2 is a known activity-dependent gene that modulates dendrite complexity (Ghiretti et al., 2014) and intrinsic excitability (Moore et al., 2018). Furthermore, the following genes were dysregulated at baseline but are not in TTX-silenced conditions: USH2A, NPAS4, MYLK2, CRH, OTOF, IGFN1, CNTN2, CPNE6 (**Figure 2E**). We extrapolate that the genes uniquely upregulated in the C9-NRE UT condition may be caused by the slight differences in excitability measured by MEA (**Figure 1**), and the genes uniquely upregulated in the C9-NRE TTX condition are negatively regulated by alterations in excitability. Genes upregulated in silenced Control i^3^Neurons (downregulated in silenced C9-NRE i^3^Neurons) relate to transcriptional regulation and the ECM, similar to in the previous comparison. The transcriptional regulators predicted to regulate genes upregulated in TTX-silenced control neurons include CGBP, TET1, DNMT1, and MLL (**Figure 2D**). Strikingly, these transcriptional regulators are related to the methylation status in cells. Moreover, CGBP binds to unmethylated CpG regions, TET1 is a DNA demethylase, DNMT1 is a DNA methyltransferase, and MLL is a histone methyltransferase. This analysis revealed synaptic genes that are consistently dysregulated in C9-NRE i^3^Neurons regardless of neuronal activity, genes that are negatively regulated by neuronal activity in C9-NRE i^3^Neurons, and genes that may play a role in the subtle results for altered network properties and synchrony phenotypes in **(Figure 1B)**.

**Figure 2.**
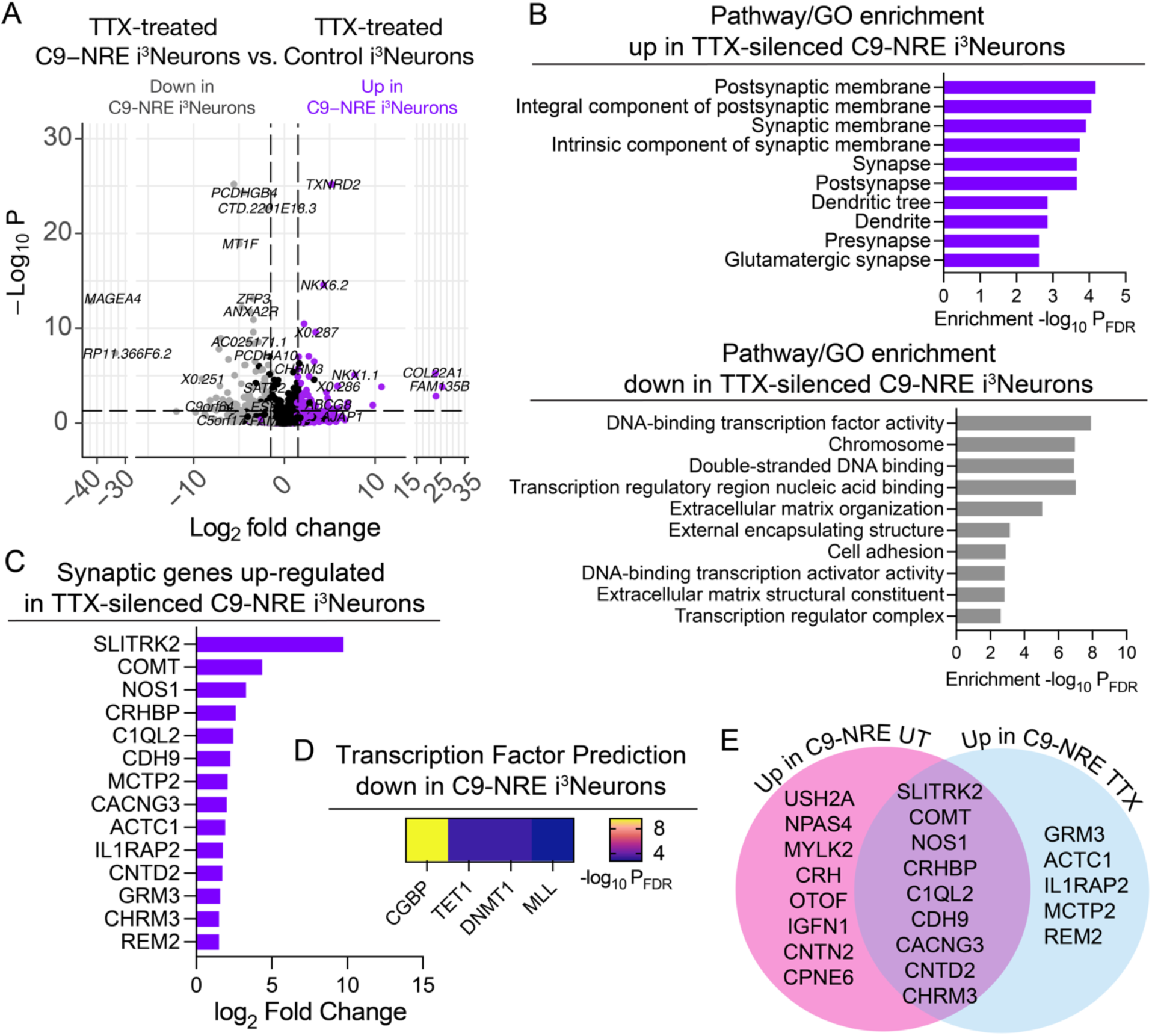
TTX-silenced C9-NRE i^3^Neurons show persistent synaptic transcript dysregulation. **(A)** Volcano plot of differentially expressed genes (DEGs) in comparing TTX-silenced Control i^3^Neurons and TTX-silenced C9-NRE i^3^Neurons. For each experiment *n* = 3 biological replicates, *m* = 2 technical replicates. **(B)** Enriched Gene Ontology (GO) terms and KEGG pathways for genes upregulated in C9-NRE and Controls are plotted. **(C)** The log_2_ fold change of synaptic genes upregulated in C9-NRE i^3^Neurons is plotted. **(D**) Predicted transcription factor activity is based on differentially expressed genes downregulated in C9-NRE i^3^Neuron. **(E)** Venn diagram of upregulated synaptic genes in the comparison of Con UT vs. C9 UT and Con TTX vs. C9 TTX.

### Neuronal depolarization induces a unique transcriptomic signature in C9-NRE i^3^Neurons

We then questioned how C9-NRE i^3^Neurons may differ in transcriptional response to controlled, increased excitation. To test this, we employed a commonly used paradigm to investigate activity-dependent transcription (Rienecker et al., 2020), which is silencing endogenous synaptic activity with TTX overnight (∼16 hours) followed by a singular depolarization event by application of high extracellular potassium chloride (55 mM KCl). In this experimental paradigm, i^3^Neurons were depolarized at time points consistent with early (2 hours KCl) and late (6 hours KCl) waves of activity-dependent gene transcription to reveal dysregulated immediate early genes and late response genes, respectively. We have previously shown that this experimental paradigm is effective at activating the activity-dependent transcription pathway in both Control and C9-NRE i^3^Neurons as measured by an increase in extracellular calcium via calcium imaging in addition to robust induction of *bona fide* immediate early genes (FOS, NPAS4), and late-response gene (BDNF) (Ghaffari et al., 2023). Importantly, we have also shown that our i^3^Neurons are resilient to this treatment even after 6 hours (Ghaffari et al., 2023). Following depolarization, we employed RNA sequencing and performed differential gene expression (**Figure 3A**). To determine how controls and C9-NRE i^3^Neurons respond to this experimental paradigm independently, we first performed differential gene expression between 0 hours of depolarization (neurons left silenced in TTX for ∼16 hours without KCl addition) and 2 hours of depolarization or TTX silenced to 6 hours of depolarization within controls or C9-NRE. While these comparisons were performed independently, the normalized expression score for all DEGs in these initial comparisons are plotted in the heat map (**Figure 3B**). There are unique gene programs that are induced selectively in C9-NRE i^3^Neurons or induced to a different degree in C9-NRE i^3^Neurons, indicating a divergence in response to this stimulus. To identify genes uniquely up- and down-regulated in C9-NRE i^3^Neurons following depolarization, we compared the DEGs identified in the control i^3^Neuron condition.

**Figure 3.**
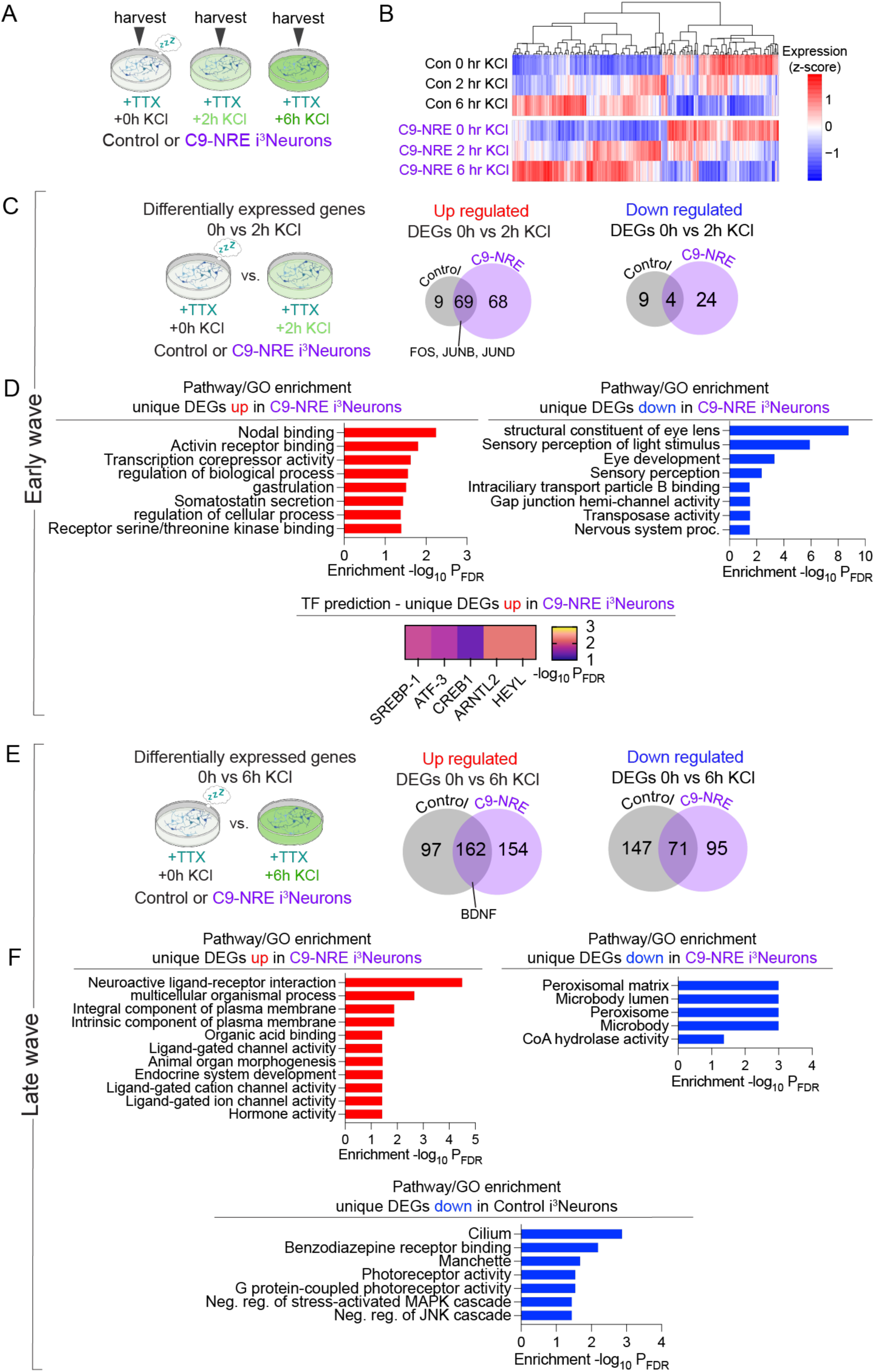
The early and late wave of activity-dependent transcription is divergent in C9-NRE i^3^Neurons. **(A)** Schema of the experimental time points of depolarization of i^3^Neurons prior to harvesting for RNA isolation. **(B)** RNA Sequencing was performed on KCl-stimulated i^3^Neurons, and differentially expressed genes (DEGs) were identified using DESeq2. Pairwise comparisons were performed between 0 hours KCl (TTX only) and 2 hours KCl or 6 hours KCl for both Control and C9-NRE i^3^Neurons. The heatmap shows the gene expression (scaled by z-score) of all DEGs between listed comparisons. **(C)** Differential gene expression for the early wave of activity-dependent gene transcription, between 0 hours KCl vs 2 hours KCl for both control or C9-NRE i^3^Neurons was performed. The number of genes uniquely up-regulated or down-regulated in control or C9-NRE i^3^Neurons are shown in the Venn diagrams. **(D)** Pathway and gene ontology (GO) analysis were performed on the uniquely up- and down-regulated genes from the comparisons in (C). Pathway analysis includes KEGG and GO analysis includes biological process, molecular function, and cellular component GO terms. The enrichment is displayed as the -log_10_ of the False Discovery Rate (FDR) for each term. **(E)** Differential gene expression for the late wave of activity-dependent gene transcription between 0 KCl vs 6 KCl for both control or C9-NRE i^3^Neurons was performed. The number of genes uniquely up-regulated or down-regulated in control or C9-NRE i^3^Neurons are shown in the Venn diagrams. **(F)** Pathway and gene ontology (GO) analysis were performed on the uniquely up- and down-regulated genes from the comparisons in (E). Pathway analysis includes KEGG and GO analysis includes biological process, molecular function, and cellular component GO terms. The enrichment is displayed as the -log10 of the False Discovery Rate (FDR) for each term.

First, we examined the early wave of activity-dependent gene transcription. Following 2 hours of depolarization, there are 68 unique genes upregulated and 24 unique genes downregulated in the C9-NRE i3Neurons (**Figure 3C**). As expected, at this time point, bona fide immediate early genes such as FOS, JUNB, and NPAS4, are upregulated in both control and C9-NRE i^3^Neurons. While levels of NPAS4 at baseline (untreated, UT) are differential between C9-NRE and Control i^3^Neurons, NPAS4 is induced at similar levels after the KCl depolarization paradigm. The genes that are uniquely upregulated in C9-NRE i^3^Neurons at 2 hours are related to Nodal binding and include CFC1 and CFC1B. The predicted regulators of these genes are CREB1, ATF-3, and SREBP-1 (**Figure 3D**), all of which are transcription factors that can be activated by neuronal activity (Zhang et al., 2011; Chen et al., 2017). Interestingly, genes uniquely downregulated in C9-NRE i^3^Neurons following 2 hours of neuronal depolarization are enriched for the many members of the crystallin family of genes, particularly of the beta and gamma subtypes. These genes include CRYBB1, CRYBB3, CRYBA2, CRYGS, CRYBA1. These results point to widespread dysregulation of the crystallin family of genes, which are not currently appreciated as genes potentially regulated by neuronal excitation.

Next, to determine if there are divergent responses in the late wave of activity-dependent gene transcription, we performed pairwise comparisons between 0 hours and 6 hours of depolarization in either control or C9-NRE i^3^Neurons (**Figure 3E**). This comparison revealed an upregulation of ligand-gated ion channels KCNK1, KCNJ16, RYR2, CHRND, in addition to plasma-membrane bound signaling molecules VSIG2, GPR50, HAS1, IL18RAP, PCDH17, APLNR, HCRTR2, SLC3A1, PTH2R, IL6R, HHIP, TNFRSF18 in C9-NRE i^3^Neurons specifically (**Figure 3F**). The cell-signaling pathways associated with these genes are varied but can be linked to cytokine-driven cell signaling. We also observed that genes associated with peroxisome regulation are downregulated in C9-NRE i^3^Neurons following 6 hours of depolarization, and these genes include NUDT7, FABP1, ACOT4, and PXT1.

Additionally, we performed pairwise comparisons between C9-NRE i^3^Neurons and Control i^3^Neurons at each time point of neuronal depolarization to account for consistent differences between genotypes (**Supplemental Figure 1A-B**). Like in prior comparisons, common themes emerged from these pair-wise comparisons. Synaptic genes were consistently upregulated in C9-NRE i^3^Neurons across all depolarization times, while genes relating to transcriptional regulation were consistently downregulated in Control (**Supplemental Figure 1C-D**). Interestingly, the crystallin family of genes was up-regulated in C9-NRE at 0 hours compared to Control i^3^Neurons at 0 hours. In previous comparisons, the crystallin family of genes was found to be downregulated upon 2 hours of depolarization. The heatmap of normalized counts of all crystallin genes in **Supplemental Figure 1E** further demonstrates the global dysregulation of this family of genes at baseline and following depolarization.

### Regression-based analysis reveals clusters of depolarization- and disease-dependent trajectory of gene programs

Due to the temporal nature of our RNA sequencing datasets, we next sought to determine if there was a model that could capture transcriptomic differences in an integrated manner and reveal novel pathways altered by depolarization in C9-NRE i^3^Neurons. Using the depolarization time (0, 2, or 6 hours) as a quantitative variable, we employed the algorithm, maSigPro (Conesa et al., 2006; Nueda et al., 2014). In this analysis, maSigPro uses regression-based analysis to identify unique gene trajectories across time and conditions. We applied this generalized linear model using the backward regression method and model-based clustering (MClust) and identified 5 distinct clusters of genes that have differential trajectories over the time of depolarization (**Figure 4A**). As in prior comparisons, the time point “0” refers to i^3^Neurons that were left silenced in TTX for 16 hours without being depolarized with KCl. The dotted line plotted in **Figure 4A** is the line of “best fit” for each cluster. It is important to note that the median profile of the trajectory for these genes across the time of depolarization is shown in **Figure 4A**, however, this is not meant to represent the trajectory of all of genes belonging to a cluster. Indeed, there are genes belonging to each cluster that are calculated to be differentially expressed in Controls and C9-NRE i^3^Neurons across time of depolarization that do not have the same trajectory direction. This can more easily be understood by looking at the expression scores for the genes in each cluster plotted in a heat map (**Supplemental A**). Cluster 1 comprises genes repressed upon depolarization in controls that are not as repressed in C9-NRE i^3^Neurons. The genes in this cluster include the transcription factor calcium-response factor (CaRF), also known as Amyotrophic lateral sclerosis 2 chromosomal region candidate gene 8 protein (ALS2CR8). ALS2CR8 or CaRF was putatively linked to ALS2 is a gene that is causative of juvenile ALS when mutated (Hadano et al., 2001). Additionally, a gene identified in Cluster 1, peroxisome proliferator-activated receptor gamma coactivator 1-alpha (PPARGC1A encoding the protein PGC-1α), that functions in mitochondrial homeostasis, is reported to be downregulated in the spinal cord of ALS patients (Thau et al., 2012), but has been shown to elicit neuroprotective effects in the SOD1 G93A mouse model (Liang et al., 2011; Zhao et al., 2011; Varghese et al., 2020). The activity-dependent transcription factor CREB1 is predicted to mediate the regulation of the genes in Cluster 1. Genes in Cluster 2 are slightly more induced in C9-NRE i^3^Neurons in response to neuronal depolarization compared to controls, although the plotted trajectory is nearly identical. Genes in Cluster 2 are enriched for cAMP signaling, modulators of neuronal excitability, and glutamatergic synapses. These genes include ATF4, CREM, SV2C, DRD2, and PICALM. Notably, DRD2, the dopamine receptor D2, has been previously identified to modulate ALS motor neuron excitability (Huang et al., 2021). Cluster 2 is predicted to be regulated by many transcription factors, but of note, are PATZ and MAZ. PATZ is a transcription factor that interacts with p53 targeting genes and p53 itself. MAZ interacts as a cofactor with CTCF1, which is predicted to regulate the genes significantly elevated in Control i^3^Neurons compared to C9-NRE neurons at baseline (**Figure 1E**). Due to these two pieces of evidence, we reason that CTCF activity is downregulated in C9-NRE i^3^Neurons compared to controls. Genes comprising Cluster 3 are more upregulated in C9-NRE i^3^Neurons following 2 and 6 hours of depolarization compared to Controls and slightly enriched for intracellular signal transduction GO terms (not shown). Cluster 3 is predicted to be regulated by activity-dependent transcription factor CREB, in addition to ETF and ATF2. Cluster 4 is repressed upon depolarization to a greater extent in C9-NRE i^3^Neurons following 2 hours of depolarization than in controls. The genes in this cluster are related to peroxisome regulation, however, they are distinct from PPARGC1A, identified in Cluster 1, and include the genes PEX12 and PEX11A. Cluster 4 enrichment terms also include GEMIN2 and GEMIN6, key components of the survival of motor neuron (SMN) complex. The transcriptional regulators predicted to regulate Cluster 4 are primarily related to methylation status and are the same regulators identified in the comparison of Control TTX vs C9-NRE TTX (**Figure 2**). These regulators include MLL, CGBP, TET1, and DNMT1. TET1 has been shown to be critical for neuronal activity-related gene expression (Rudenko et al., 2013). Cluster 5 is comprised of genes always expressed lower in C9-NRE i^3^Neurons regardless of depolarization compared to Controls, but also differ in the trajectory across depolarization. Genes related to kinase activity are the primary drivers of this cluster. These genes include PPP1R1B, CCNB3, and HEXIM2. Cluster 5 is predicted to be regulated by the aptly named transcriptional regulator Yin Yang 1 (YY1); its role depends on protein partners’ availability to modulate the activity (He and Casaccia-Bonnefil, 2008) and can act both as a transcriptional activator and repressor (Verheul et al., 2020). YY1 has critical neuron-specific functions in cortical excitatory neurons and dysregulation of YY1 has been linked to neurodevelopmental and neurodegenerative diseases (Pabian-Jewula et al., 2022). It has been suggested that YY1 plays a role in repressing transcription of the glutamate transporter EAAT2 (Pabian-Jewula et al., 2022), a loss of which is linked to ALS (Rosenblum and Trotti, 2017). Collectively, this analysis revealed that pathways pertaining to genome methylation, modulation of neuronal excitability, and peroxisomal regulation are dysregulated differentially in Control and C9-NRE i^3^Neurons following membrane depolarization.

**Figure 4.**
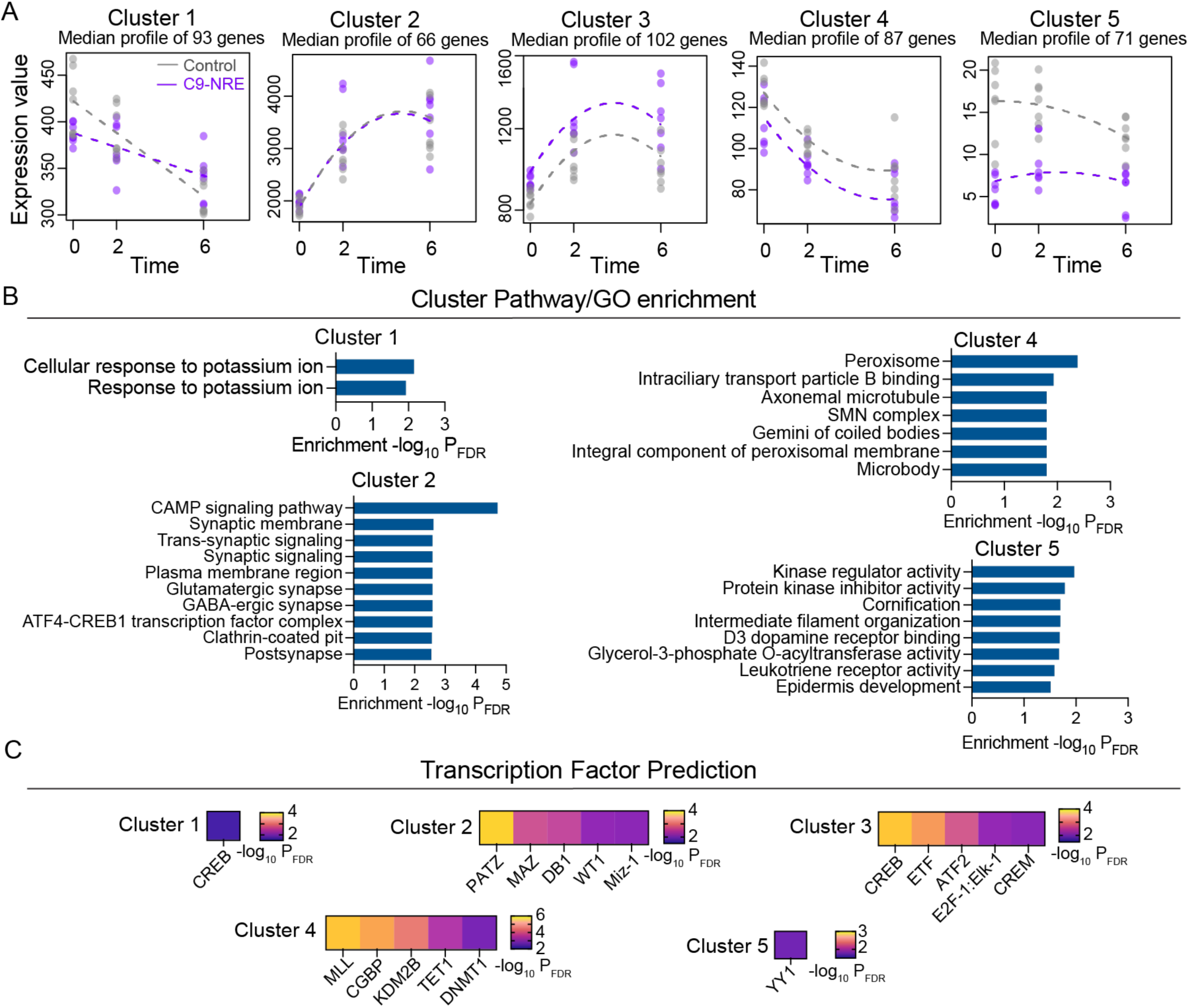
Profiling temporal dynamic of gene expression following neuronal depolarization captures distinct transcriptomic features dysregulated in C9-NRE i^3^Neurons. **(A)** DEG clusters were identified using maSigPro, a regression-based approach to find genes for significant gene expression profile differences between experimental groups in time course RNA-Seq experiments. **(B)** Pathway Enrichment/GO for each cluster identified via maSigPro. **(C)** Transcription factor prediction is based on the differentially expressed genes in each cluster.

### The transcriptome of modeling the excitability phenoconversion in C9-ALS/FTD

To determine how the activity-dependent transcriptome in C9-NRE i^3^Neurons may be shaped by neuronal activity more closely mimicking cortical hyperexcitability, we used the voltage-gated potassium channel blocker tetraethylammonium (TEA). It has been reported that the excitability of C9-ALS/FD motor neuron iPSC models show initial hyperexcitability and then transitions to hypoexcitability once cultured further (Sareen et al., 2013; Devlin et al., 2015; Burley et al., 2022), which mimics what is observed in patients; initial cortical hyperexcitability to eventual loss of neuronal activity with neurodegeneration. Application of TEA led to an elevation in spike number of both Control and C9-NRE i^3^Neurons as measured by an increase in peak amplitude and frequency, measured by fluorescence intensity of the calcium indicator Fluo-4 and the MATLAB program PeakCaller (Artimovich et al., 2017) (**Figure 5A**). The calculated area under the curve (AUC) shows a similar level of response to TEA in both control and C9-NRE i^3^Neurons. i^3^Neurons were incubated with TEA for 24 hours prior to harvesting and submission to RNA seq. To complement the hyperexcitation paradigm, and to model the phenoconversion from hyperexcitability to hypoexcitability, we silenced endogenous synaptic activity with TTX for 24 hours (data previously shown in the pairwise comparisons in **Figure 1**). While silencing endogenous activity with TTX does not mimic the full complement of hypoexcitability, this approach ensures high signal-to-noise ratio for transcriptomic readouts. We first performed pairwise comparisons between UT Control i^3^Neurons and TTX-silenced Control i^3^Neurons, which did not yield significant results, A similar result was observed in the comparison between UT C9-NRE i^3^Neurons and TTX-silenced C9-NRe i^3^Neurons. We reasoned that, due to the varied nature of firing patterns of individual i^3^Neurons *in vitro,* the signal-to-noise ratio might be too low to detect significant activity-dependent gene differences in bulk transcriptomic sequencing. We then compared UT versus TEA and TTX versus TEA for both Control and C9-NRE i^3^Neurons (**Supplemental Figure 2**). We found shared and unique DEGs for these comparisons in Control and C9-NRE i^3^Neurons. Notable results include the peroxisomal gene, PPARGC1A is uniquely up in C9-NRE i^3^Neurons in the UT versus TEA comparison and TTX versus TEA comparison. In prior comparisons using the KCl paradigm and regression analysis, PPARGC1A was found to be a member of Cluster 1 (**Figure 4A**), indicating a differential trajectory between Control and C9-NRE i^3^Neurons across the time of depolarization. These findings suggest a convergence of neuronal excitation gene expression in C9-NRE neurons (**Supplemental Figure 2A,C**). The voltage-gated calcium channel CACNA1G was found to be uniquely upregulated in C9-NRE i^3^Neurons in the UT vs. TEA and TTX vs. TEA comparisons, comprising the “low voltage-gated calcium channel activity” GO term (**Supplemental Figure 2A**). The GO term “reg. of hormone levels”, uniquely upregulated in C9-NRE i^3^Neurons in the TTX vs TEA comparison, includes the neurotrophin NGF (nerve growth factor) and the neuropeptide precursor VGF. Overall, while there is a level of convergence between the two experimental paradigms, there are genes induced by TEA stimulation that were not previously identified to be induced by KCl depolarization.

**Figure 5.**
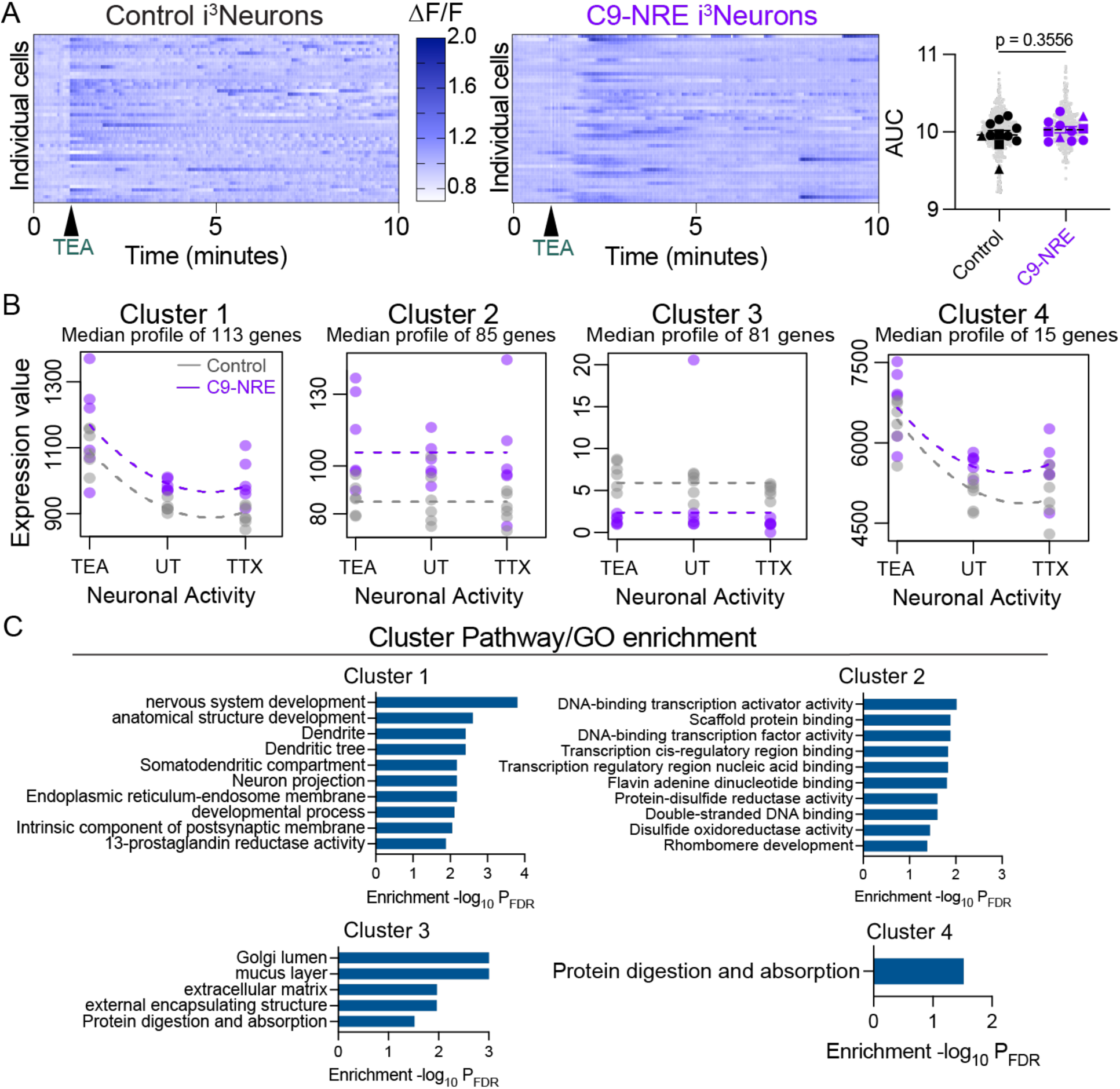
Modeling the C9-NRE patient hyper- to hypo-excitability spectrum using i^3^Neurons identifies disease-relevant pathways. **(A)** A representative raster plot of i^3^Neurons loaded with Ca^2+^ indicator dye Fluo-4 and, following a baseline recording, were stimulated with 4 mM TEA as shown by the arrow at 1 minute. The ΔF/F was calculated per cell using PeakCaller and is plotted. *n* = 3 biological replicates, *m* = 3-4 independent experiments per line, *o* = 50 cells per experiment. **(B)** DEG clusters were identified using maSigPro, using levels of neuronal activity as pseudo-time. **(C)** Pathway Enrichment/GO for transcripts divergently regulated in C9-NRE i^3^Neurons compared to Control i^3^Neurons following silencing with TTX or activation with TEA.

We then considered each experimental paradigm as a state of excitability that mimics the dimension of the disease state across time (pseudotime) from hyperexcitability (TEA treatment) to hypoexcitability (TTX treatment). Using this conceptual framework, we applied maSigPro regression analyses to this dataset to determine genes that have statistically significantly different trajectories across “disease” pseudotime in Control versus C9-NRE i^3^Neurons. Using model-based clustering, we identified 4 distinct clusters of significantly different genes (**Figure 5B-C**). Cluster 1 is enriched for genes more sensitive to changes in activity in C9-NRE i^3^Neurons. It appears there is a Goldilocks level of neuronal activity for this cluster of genes; upregulated with TEA treatment and TTX, but consistently higher in C9-NRE i^3^Neurons regardless of treatment. Interestingly, these genes are enriched for modulators of excitability and neuron-specific compartments (**Figure 5D**), perhaps indicating an attempt at neuronal homeostasis in regard to excitability. These synaptic genes include CACNG3, COMT, and DRD2. CACNG3 and COMT were previously identified to be dysregulated in untreated and TTX-silenced C9-NRE i^3^Neurons (**Figure 2D**), and DRD2 was also dysregulated by neuronal depolarization in Cluster 2 (**Figure 4A**). Interestingly, Cluster 4 has a similar trajectory to Cluster 2, however, it differs in magnitude of expression levels, and includes the synaptic genes CNTN2, and RAB3C. Cluster 1 is enriched for protocadherin genes PCDHA10, PCDHGB4, PCDH19. Protocadherins are cell adhesion molecules that are highly important in CNS functions such as circuit formation and neuronal survival (Pancho et al., 2020). Additionally, mutations in PCDH19 are causative of epilepsy and neurodevelopmental diseases (Dell’Isola et al., 2022). Cluster 2 genes that are consistently up in C9-NRE i^3^Neurons compared to Controls and seem to be increased when silenced, including NOS1 and CDH9, regardless of neuronal activity. There is significant enrichment for GO terms related to transcriptional machinery, which may promote the diverse transcriptional response in C9-NRE i^3^Neurons. Cluster 3 is enriched for extracellular matrix GO terms, and Cluster 4 is enriched for the single GO term ‘protein digestion and absorption’, which includes the genes SLC38A2 and ATP1B1. Collectively, our coupling of transcriptomics to experimentally modeling the phenoconversion of hyperexcitability to hypoexcitability yields novel insights into gene profiles significantly altered in Control and C9-NRE i^3^Neurons.

## Discussion

In this work, we show that at baseline, C9-NRE patient-derived iPSCs differentiated into cortical-like i^3^Neurons show mild alterations in synaptic gene expression and firing properties when compared to healthy control i^3^Neurons, similar to findings in iPSC motor neurons (Sareen et al., 2013; Ababneh et al., 2020). Furthermore, following experimental conditions that mimic increased neuronal activity, we show that membrane depolarization or blockade of potassium channels induce a unique and specific transcriptomic signature in C9-NRE i^3^Neurons. Strikingly, in many of our analyses, genes involved in transcriptional machinery were downregulated in C9-NRE i^3^Neurons. We hypothesize that, in concert with neuronal activity, downregulated transcriptional genes prime C9-NRE i^3^Neurons for dysfunctional transcriptional responses to neuronal excitation. The precise influence of downregulated transcriptional genes on dysregulated synaptic genes at baseline, or vice-versa, was not determined in this study but systematically manipulating genes in these pathways may reveal the causal order in which these events occur. However, it seems most likely that intrinsic differences in gene networks in C9-NRE carriers could serve as primer for synaptic alterations. Many of these synaptic genes that are consistently upregulated in C9-NRE i^3^Neurons in many comparisons include, NOS1, CDH9, and CACNG3 (**Figure 1E**, **Figure 2C**, **Supplemental Figure 1C**). These results mimic a recent proteomics study in which NOS1, CDH9, and CACNG3 were more highly enriched in synaptosomes isolated from pre-frontal cortices of C9-NRE carriers (Laszlo et al., 2022). Strikingly, all 3 of these genes are targets of the known activity-dependent transcription factor NPAS4, which plays a role in the balance of excitatory-inhibitory neurotransmission (Fu et al., 2020). NPAS4 has recently been found to be a target of the Alzheimer’s disease-relevant protein amyloid precursor protein (APP) and is lost with ablation of APP (Opsomer et al., 2020). Collectively, work linking NPAS4 to a neurodegenerative disease-relevant protein, the established role of NPAS4, and our data implicate NPAS4 in physiology and disease states. Our data also shows dysregulation of the gene SLITRK2, which has been found to be similarly upregulated in C9-NRE patient tissue and iPSC motor neurons (Sareen et al., 2013; Satoh et al., 2014; Barbier et al., 2021). Strikingly, SLITRK2 upregulation in C9-NRE FTD frontal cortex is associated with a SNP near SLITRK2, which is associated with an earlier onset of dementia (Barbier et al., 2021). These results highlight the relevance of our findings in cortical-like i^3^Neurons. Dysfunction of the activity-dependent gene transcription pathway can manifest into neurological disease (Roussos et al., 2016; Osenberg et al., 2018; Boulting et al., 2021).

Cortical hyperexcitability is a relevant event in the early stages of ALS disease progression in patients (Vucic et al., 2008; Kiernan, 2009; Bae et al., 2013; Vucic et al., 2013; Menon et al., 2020; Vucic et al., 2021; Shibuya et al., 2022), but neither the cause nor consequences are well understood. In this work, we sought to understand the downstream effects of altered neuronal excitation by investigating gene expression in a ALS/FTD-relevant disease model system, C9-NRE cortical neurons following two modes of neuronal stimulation. The first mode of neuronal stimulation we utilized was membrane depolarization. While this is not a mimic of hyperexcitability, it is well-understood, and therefore, we were able to validate that our system responded in an established fashion. The second mode of neuronal stimulation used in this study was blockade of potassium channels with tetraethylammonium (TEA), which is a mimic of hyperexcitability. Exposure of i^3^Neurons to TEA yields unique results compared to KCl depolarization, indicative of mode-specific results. Using time as a quantitative variable, we employed regression analysis that revealed striking patterns of the trajectory of gene expression across time of depolarization in the KCl paradigm and ‘pseudotime’ in the TEA paradigm. These analyses of KCl-stimulated i^3^Neurons revealed there are significant shifts in immediate early genes and late-response genes. For this analysis, the plotted trajectory for a given Cluster generated by the regression analysis (maSigPro (Conesa et al., 2006; Nueda et al., 2014)) indicates the median profile of the genes identified to be differentially regulated across time. However, not all genes in each cluster have this precise trajectory. These results on stimulated healthy and C9-NRE i^3^Neurons reveal novel and established dysregulated pathways in ALS, emphasizing the importance of neuronal activity to pathogenic pathways. Among the established pathways involved in ALS are peroxisomal regulation (Liang et al., 2011; Zhao et al., 2011; Thau et al., 2012; Varghese et al., 2020), dysregulation of players in the survival motor neuron (SMN) complex , calcium-response factor (CaRF), also known as Amyotrophic lateral sclerosis 2 chromosomal region candidate gene 8 protein (ALS2CR8) (Hadano et al., 2001), and genome-wide methylation status (Ebbert et al., 2017; Zhang et al., 2017; Jackson et al., 2020). Genes involved in peroxisomal regulation were identified to be dysregulated in many of our analyses. The gene PPARGC1A (peroxisome proliferator-activated receptor gamma coactivator 1-alpha), which encodes the protein PGC-1α, functions to maintain mitochondrial homeostasis and biogenesis, is downregulated in ALS patient spinal cord (Thau et al., 2012). Interestingly, studies in which ectopic neuronal overexpression of PGC-1α in the SOD1 G93A mouse model led to a rescue of motor performance, survival, and restored dysfunctional mitochondrial activities (Liang et al., 2011; Zhao et al., 2011; Varghese et al., 2020). PPARGC1A was identified to be a member of Cluster 1 in the regression analysis of the KCl experimental paradigm, indicating a differential trajectory over time of membrane depolarization. Although belonging to Cluster 1 of the regression model of the KCl paradigm, PPARGC1A is upregulated in response to membrane depolarization in Control and C9-NRE i^3^Neurons, although with a different trajectory (**Supplemental C**). Interestingly, PPARGC1A is also uniquely upregulated in C9-NRE i^3^Neurons in the UT versus TEA pairwise comparison and TTX versus TEA pairwise comparison. However, when looking carefully, PPARGC1A is also induced by neuronal excitation in Control i^3^Neurons in the UT versus TEA pairwise comparison and TTX versus TEA pairwise comparison, however, slightly below the log2 fold change cutoff of 1.5, and therefore was excluded in our analyses. Other peroxisomal genes found to be dysregulated include the PPARGC1A targets PEX12 and PEX11A identified as members of Cluster 4 in the KCl paradigm regression analysis (**Figure 4A**). Peroxisomal genes were also identified to be dysregulated in the pairwise comparisons 0 hours KCl versus 6 hours KCl in C9-NRE i^3^Neurons (**Figure 3F**), and some of these genes include are PPARGC1A targets; NUDT7, FABP1, ACOT4, and PXT1. Methylation status may also be differentially altered in C9-NRE i3Neurons. Based on the predicted reduced activity of TET1 in C9-NRE i^3^Neurons it may be that some TET1 targets are hypermethylated and therefore cannot be transcribed. DNMT1 regulates neuronal activity levels (Bachmann et al., 2021), and mediates age-related degeneration via proteostatic imbalance (Hahn et al., 2020). Furthermore, loss of DNMT1 itself, and therefore hypomethylation, in excitatory neurons leads to impaired excitability (Hutnick et al., 2009). DNA methylation status, which is regulated by neuronal activity , has been heavily implicated in C9-ALS and sALS (Ebbert et al., 2017; Zhang et al., 2017). Overall, a major theme emerging from Cluster 4 is genes that are more heavily repressed in C9-NRE i^3^Neurons may be methylation-status dependent, which is similar to our earlier findings (**Figure 2**). Loss of methylation or demethylation has widespread genomic consequences, which may prime neurons to degenerative pathways.

The novel pathways revealed by the membrane depolarization paradigm include widespread Crystallin gene dysregulation. Among the genes uniquely downregulated in C9-NRE i^3^Neurons following depolarization are the beta and gamma subtypes of the crystallin family of genes, including CRYBB1, CRYBB3, CRYBA2, CRYGS, and CRYBA1. The crystallin family of genes are primarily responsible for the maintenance of the eye lens, but also have a role in anti-apoptotic signaling and neuroprotection (Mao et al., 2004; Liu et al., 2022). However, these reports have been primarily using retinal models, and so the role of crystallin genes in cortical-like neurons remains to be understood (Graw, 2009, 2017). Furthermore, the alpha subtype of crystallin genes have been implicated in the presence of neurofibrillary tangles in Alzheimer’s disease (Mao et al., 2001). The significant repression of these genes upon depolarization could lead to neuronal vulnerability, perhaps priming susceptibility to a second hit.

Experimentally, we modeled the phenoconversion of hyperexcitability to hypoexcitability by using TEA and TTX as “disease states” indicative of the status of neuronal activity. By attributing the level of neuronal activity to time-of-disease, with high neuronal activity as early in the disease, and low neuronal activity as late in the disease, we were able to use this “pseudotime” as a quantitative variable and employed the regression-based analysis, maSigPro (Conesa et al., 2006; Nueda et al., 2014) to determine clusters of genes that are dysregulated across psuedotime. We are the first to report transcriptomic alterations in concert with disease-relevant state of neuronal activity in the context of C9orf72-linked ALS/FTD. This analysis revealed dysregulated pathways similar to the KCl paradigm, indicating a convergent response to neuronal excitation. As previously mentioned, the peroxisomal pathways were similarly altered in C9-NRE i^3^Neurons using KCl depolarization and TEA stimulation. Additionally, several protocadherin genes were identified to be dysregulated by TEA stimulation. Interestingly, one of the protocadherin genes, PCDH19, is linked to epilepsy . Another gene upregulated in C9-NRE i^3^Neurons following TEA stimulation is CACNA1G, which is also linked to the epilepsy (Berecki et al., 2020). It is a testament to our experimental paradigm which aimed to mimic a hyperexcitable state, transitioning to hypoexcitable state, that genes required for normal excitability to be disrupted in this condition.

Collectively, our data shows baseline synaptic dysfunction and altered responses to neuronal excitation in C9-NRE i^3^Neurons. The next logical inquiry is the reasoning for why these altered responses occur. There are several reasons, which relate to the three proposed pathogenic mechanisms of the C9-NRE. Firstly, the loss of C9orf72 protein may impact how neurons respond to stimuli. C9orf72 can be localized to lysosomes (Amick et al., 2016; Sullivan et al., 2016; Laflamme et al., 2019; Lall et al., 2021) and the synapse (Frick et al., 2018; Xiao et al., 2019; Bauer et al., 2022), and we reason that because these two compartments respond to the needs of the cell, a loss of C9orf72 may impact this response. It could also be the gain-of-function mechanisms, repeat-containing RNA, or dipeptide repeat proteins, that mediate either the dysfunction observed at baseline or the altered response to neuronal excitation. Repeat-containing RNA is known to sequester RNA-binding proteins and impair proper RNA metabolism , which may lead to an improper elevation or degradation of transcripts. Furthermore, the dipeptide repeat proteins (DPRs) that are translated can impact the neuronal response to excitation and vice versa (Westergard et al., 2019; Jensen et al., 2020). A worthy avenue of pursuit would be to understand how the 3 pathogenic mechanisms stemming from the C9-NRE impact the responses we have observed in our analyses. This could be done by correlating RNA foci and DPR levels and subcellular localization with our transcriptomic data. Furthermore, the replacement of C9orf72 in i^3^Neurons followed by the experimental paradigms we employed would elucidate how the loss of C9orf72 protein may have impacted these responses to excitation. It will be critical to determine whether any alterations we observe *in vitro* confer resilience or vulnerability. Additionally, future studies to determine the precise role of the predicted transcription factors in these identified altered pathways are necessary.

## Materials and Methods

### hiPSC culturing and i^3^Neuron differentiation

hiPSC-i^3^Neuron lines were generously gifted by Dr. Sami Barmada (University of Michigan) (Weskamp et al., 2020), and Dr. Michael Ward (NIH) (Fernandopulle et al., 2018). Two healthy control and two C9-NRE carrier hiPSC-i^3^Neuron lines were obtained from Dr. Barmada, and one healthy control and one C9-NRE carrier hiPSC-i3Neuron line were obtained from Dr. Michael Ward (NIH). The patient demographic information for each line is provided in Table S1. All hiPSCs were maintained in Essential 8 Flex media (Gibco) on Matrigel-coated plates (Corning). Differentiation was performed using the techniques outlined in (Fernandopulle et al., 2018). Briefly, to differentiate iPS cells into cortical neurons, cells were dissociated into single cells and plated on Matrigel-coated plates in induction media (DMEM/F12, N2, NEAA, and L-glutamine) containing 2 mg/ml doxycycline. Cells were induced into neurons by treatment with doxycycline for 3 days before being dissociated and plated on PLO-coated plates. Cells were then allowed to mature for an additional 22 days in cortical maturation media consisting of BrainPhys media (STEMCELL) supplemented with BDNF (Peprotech), NT-3 (Peprotech), B27 (Gibco), and Laminin (Thermo Fisher), with half media changes twice weekly. Neurons were harvested or experimented upon at 25 days post-differentiation (3 days of induction, 22 days of maturation).

### Stimulation of i^3^Neurons

For KCl depolarization experiments: i^3^Neurons were silenced overnight (16 hours) with 1 µM TTX. The following day, neurons were stimulated with KCl depolarization buffer (170 mM KCl, 2 mM CaCl_2_, 1 mM MgCl_2_, 10 mM HEPES, solution pH 7.4) as previously described (Kim et al., 2010), or with an equal volume of media (vehicle treated). Neurons were depolarized for 0 hours (vehicle), 2 hours, or 6 hours and were harvested at each time point. For TEA stimulation experiments: tetraethylammonium (TEA) was added to culture media for a final concentration of 4 mM TEA or with a volume-matched vehicle control (media). Neurons were stimulated for 0 hours (vehicle), 2 hours, or 6 hours.

### Calcium imaging

For calcium imaging experiments, i^3^Neurons were washed with aCSF (10 mM HEPES, 1 mM MgCl_2_, 2 mM CaCl_2_, 10 mM glucose, 145 mM NaCl, 5 mM KCl), and loaded with 1 µM Fluo-4 AM for 15 minutes. Neurons were then washed twice with aCSF and incubated for 5 minutes in between each wash prior to the start of recording. Neurons were imaged on a confocal microscope (Nikon A1R) with a 10x objective every 3 seconds for 10 minutes. A baseline recording of 1 minute was captured prior to perfusion of either high-KCl containing buffer (10 mM HEPES, 1 mM MgCl_2_, 2 mM CaCl_2_, 10 mM glucose, 95 mM NaCl, 55 mM KCl) or of TEA-containing buffer (aCSF + 4 mM TEA). Regions of interest (ROIs) were captured using NIS Elements software. Normalized fluorescence values over time (ΔF/F) was calculated and the area under the curve (AUC) was calculated in GraphPad Prism 9. For TEA experiments, ΔF/F was calculated using the Matlab GUI PeakCaller [70] with the following parameters: required rise = 15% relative, max lookback = 25, required fall = 15% relative, max lookahead = 25, interpolate across closed shutters = true, trend control = exponential moving average (1-sided), trend smoothness = 60.

### RNA sequencing and bioinformatic analysis

RNA was isolated using RNeasy Plus Mini Kit (Qiagen), and cDNA was synthesized from 500 ng RNA using a QuantiTect Reverse Transcription Kit (Qiagen). Library preparation and RNA sequencing were performed by Novogene and subjected to QC analysis according to their established protocols. Clean .fastq files were aligned to the hg38 genome using STAR (v.2.7.10b) (Dobin et al., 2013). Read summarization was performed on .bam files to obtain a featurecounts file using the R package Rsubread (v.2.12.3) (Liao et al., 2019). The feature counts file was then used as input for Differential Expression analysis using the R-package DESeq2 (v.1.36.0) (Love et al., 2014). A log_2_ fold-change cutoff of 1.5 and a p-value of less than or equal to 0.05 were used to determine the significance of differentially expressed genes. In analyses where maSigPro (Conesa et al., 2006; Nueda et al., 2014) was used, a “backwards” step method was used. The online tools ShinyGO and gProfiler were used for gene ontology and KEGG pathway enrichments. Redundant terms were discarded using REVIGO or manual filtering. SynGO was used to determine the synaptic transcripts enriched in comparisons in Figure 1 and Figure 2. For main figures, no more than the top 10 GO or KEGG terms, and the top 5 predicted regulators were shown. All raw RNA sequencing files are available online (GSE275765) and this dataset has been made publicly available for exploration in a Shiny application (https://haeuslerlab.shinyapps.io/ltg_shiny/).

### Multi-electrode Array (MEA)

Cortical-like i^3^Neurons were plated on a CytoView MEA plate (Axion Biosystems) at day 3 of differentiation. MEA recordings were performed on a Maestro Edge (Axion Biosystems) every 3-4 days during differentiation. Recordings on the MEA were performed prior to media changes. Neurons were removed from the incubator and were allowed to equilibrate in the recording chamber for 15 minutes prior to the start of recording. Neurons were recorded for 15 minutes, and the metrics shown in this study are from 15-minute recordings.

### Statistical analysis

All statistical analyses on data, excluding RNA-sequencing analyses, were performed using GraphPad Prism 9 software, as described in each Figure legend. Statistical analysis of RNA-sequencing is described above.

## Acknowledgments / Funding Sources

We would like to thank the Jefferson Weinberg ALS Center members for providing valuable feedback on this manuscript. We would also like to acknowledge Sami Barmada and Michael Ward for generously gifting us with i^3^Neuron lines. Additional figures were created with BioRender.com. This work was supported by the following sources: RF1NS114128 and R01NS114128 to A.R.H; R01NS109150 to P.P.; the Farber Family Foundation and the Aldrich Foundation to the Jefferson Weinberg ALS Center.

## Conflict of Interest

The authors declare no conflicts of interest.

## Author Contributions

LTG and ARH conceptualized the research with support from PP and DT. LTG designed all experiments with input from ARH. LTG performed all experiments except; EW performed alignment of fastq files and read summarization for RNA sequencing analysis. LTG, AB, and KC performed MEA experiments. All data analyses were overseen by ARH. LTG led the manuscript writing along with ARH. LTG and ARH developed and assembled the figures. PP, DT provided comments and edits to the manuscript. All authors read and approved the final manuscript.

## Declaration of generative AI and AI-assisted technologies in the writing process

During the preparation of this work the author (s) used ChatGPT and/or Grammarly in order to improve grammar and clarity. After using this tool/service, the author (s) reviewed and edited the content as needed and take (s) full responsibility for the content of the publication.

---

**Supplemental Figure 1.**
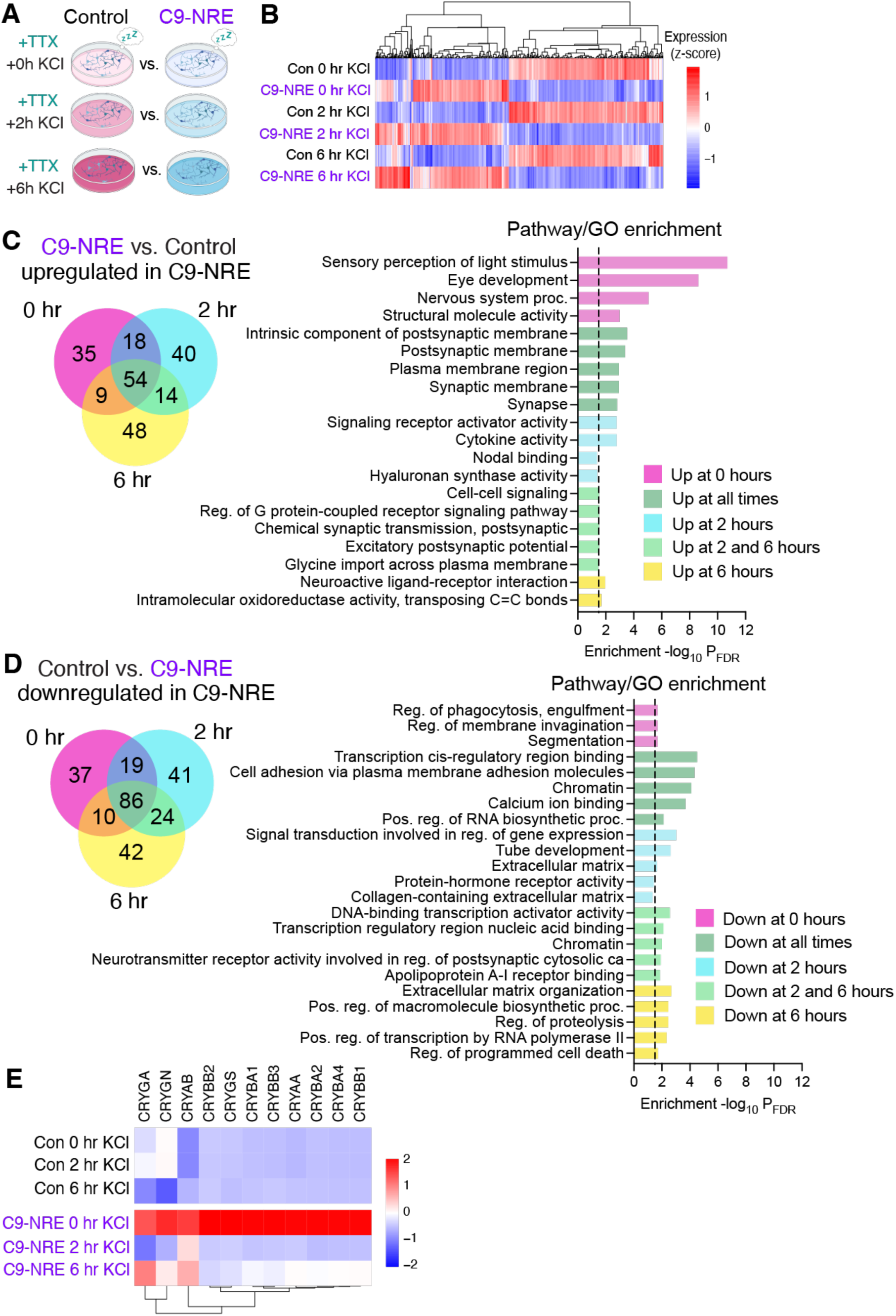
Pairwise comparisons show C9-NRE i^3^Neurons are enriched for receptor-signaling genes and are deficient in extracellular matrix-signaling pathways upon depolarization. **(A)** Schema of origin of DEGs from DESeq2 used in Venn diagram comparisons. At each time point (0, 2, 6), Control i^3^Neurons are compared to C9-NRE i^3^Neurons. **(B)** Hierarchical clustering heatmap showing the gene expression (scaled by z-score) of pooled DEGs between listed comparisons in (A). (**C**) Venn diagram of DEGs shows the inherent transcriptomic differences between Control and C9-NRE i^3^Neurons and how depolarization drives differential gene expression. Pathway Enrichment/GO are for transcripts upregulated in C9-NRE i^3^Neurons. (**D**) same as in (B) but for down-regulated transcripts in C9-NRE i^3^Neurons. (**E**) Hierarchical clustering heatmap showing widespread dysregulation of the crystallin family of genes in C9-NRE i^3^Neurons.

**Supplemental Figure 2.**
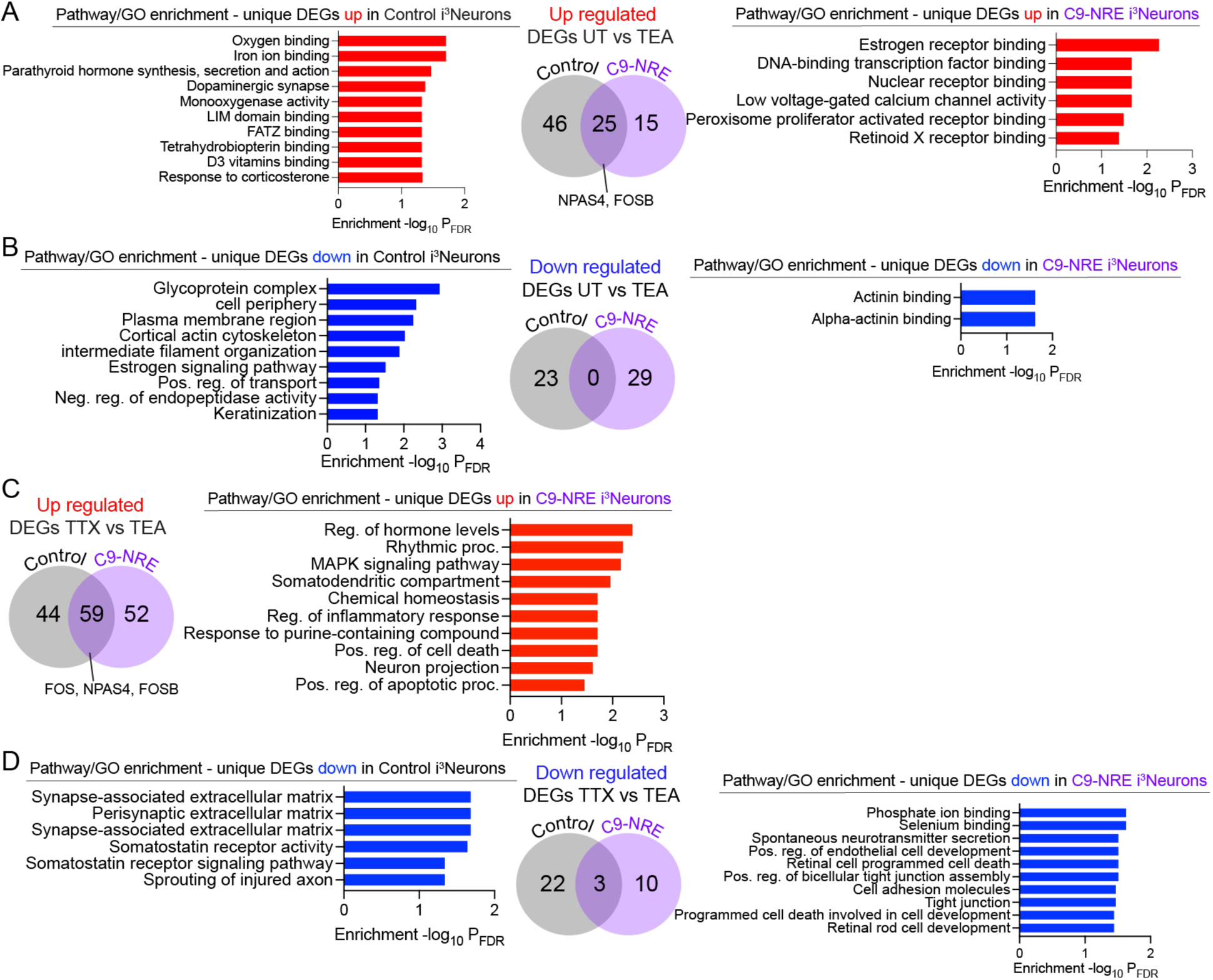
Pairwise differential gene expression of TEA-stimulated and TTX-silenced i^3^Neurons. RNA Sequencing was performed on TTX-silenced and TEA-stimulated i^3^Neurons, and differentially expressed genes (DEGs) were identified using DESeq2. Pairwise comparisons were performed between untreated (UT) and TEA stimulated, or TTX-silenced and TEA stimulated for both Control and C9-NRE i^3^Neurons. The number of genes uniquely up-regulated or down-regulated in control or C9-NRE i^3^Neurons are shown in the Venn diagrams. **(A)** Pathway and gene ontology (GO) analysis were performed on the uniquely up-regulated DEGs in either Control or C9-NRE i^3^Neurons in UT vs. TEA **(B)** Same as in (A) but for down-regulated genes. **(C)** Pathway and gene ontology (GO) analysis were performed on the uniquely up-regulated DEGs in either Control or C9-NRE i^3^Neurons in TTX vs. TEA. **(D)** Same as in (C) but for down-regulated genes. Pathway analysis includes KEGG and GO analysis includes biological process, molecular function, and cellular component GO terms. The enrichment is displayed as the -log_10_ of the False Discovery Rate (FDR) for each term.

**Supplemental Figure 3.**
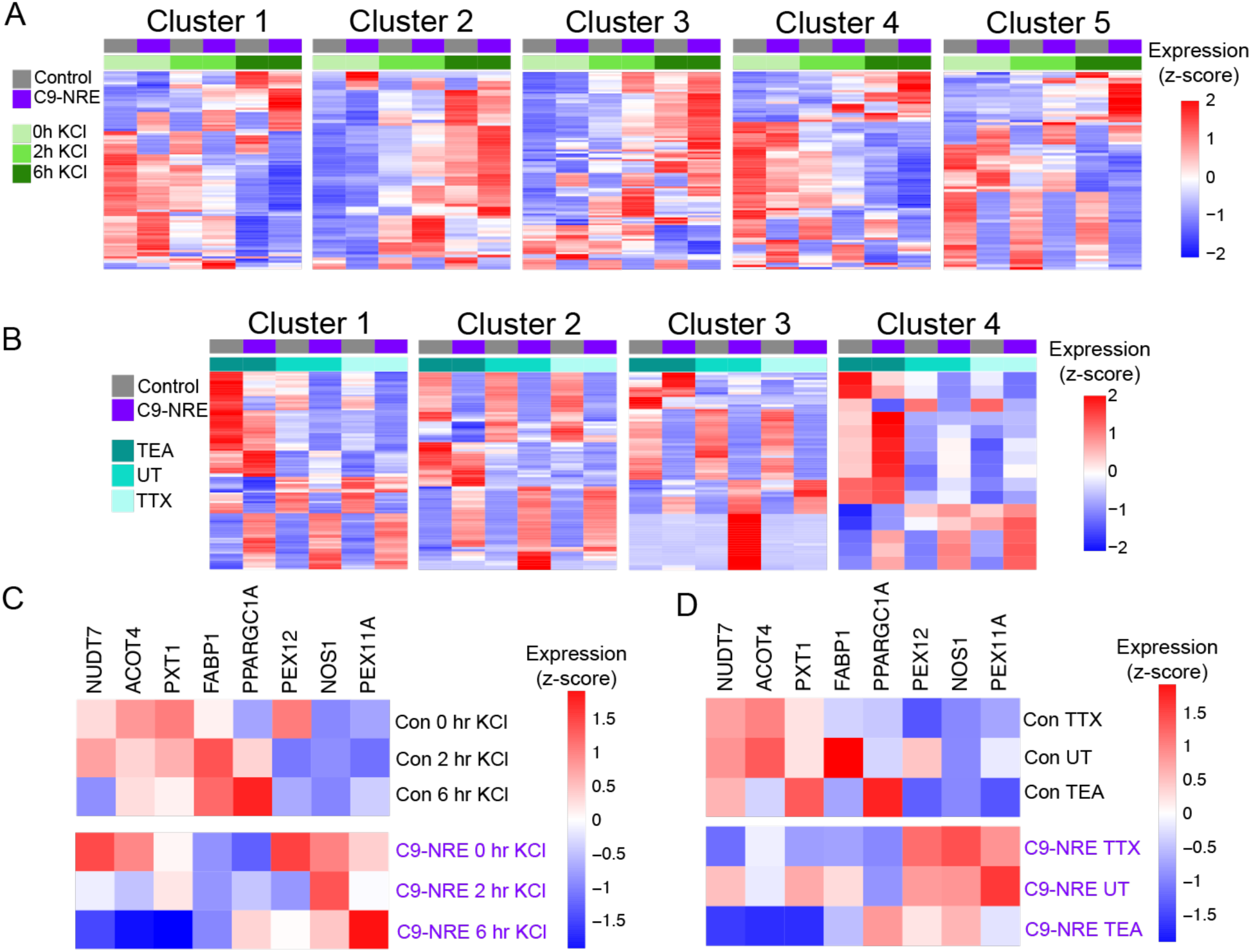
Heatmaps of Clusters identified by maSigPro and peroxisomal gene dysregulation. **(A)** DEG clusters were identified using maSigPro, using levels of neuronal activity as pseudo-time. **(B)** DEG clusters were identified using maSigPro, using levels of neuronal activity as pseudo-time with TEA stimulation, UT (untreated), and TTX silenced i^3^Neurons. **(C)** A heatmap showing the dysregulation of peroxisome-associated genes in Control and C9-NRE i^3^Neuron samples used in the KCl paradigm. **(D)** A heatmap showing the dysregulation of peroxisome-associated genes in Control and C9-NRE i^3^Neuron samples used in the TEA paradigm.

## Notes

https://www.ncbi.nlm.nih.gov/geo/query/acc.cgi?acc=GSE275765

### Competing Interest Statement

The authors have declared no competing interest.

## References

1. Ababneh NA, Scaber J, Flynn R, Douglas A, Barbagallo P, Candalija A, Turner MR, Sims D, Dafinca R, Cowley SA, Talbot K (2020) Correction of amyotrophic lateral sclerosis related phenotypes in induced pluripotent stem cell-derived motor neurons carrying a hexanucleotide expansion mutation in C9orf72 by CRISPR/Cas9 genome editing using homology-directed repair. Hum Mol Genet 29:2200–2217.

2. Amick J, Roczniak-Ferguson A, Ferguson SM (2016) C9orf72 binds SMCR8, localizes to lysosomes, and regulates mTORC1 signaling. Mol Biol Cell 27:3040–3051.

3. Artimovich E, Jackson RK, Kilander MBC, Lin YC, Nestor MW (2017) PeakCaller: an automated graphical interface for the quantification of intracellular calcium obtained by high-content screening. BMC Neurosci 18:72.

4. Bachmann S, Linde J, Bell M, Spehr M, Zempel H, Zimmer-Bensch G (2021) DNA Methyltransferase 1 (DNMT1) Shapes Neuronal Activity of Human iPSC-Derived Glutamatergic Cortical Neurons. Int J Mol Sci 22.

5. Bae JS, Simon NG, Menon P, Vucic S, Kiernan MC (2013) The puzzling case of hyperexcitability in amyotrophic lateral sclerosis. J Clin Neurol 9:65–74.

6. Barbier M et al. (2021) SLITRK2, an X-linked modifier of the age at onset in C9orf72 frontotemporal lobar degeneration. Brain 144:2798–2811.

7. Bauer CS, Cohen RN, Sironi F, Livesey MR, Gillingwater TH, Highley JR, Fillingham DJ, Coldicott I, Smith EF, Gibson YB, Webster CP, Grierson AJ, Bendotti C, De Vos KJ (2022) An interaction between synapsin and C9orf72 regulates excitatory synapses and is impaired in ALS/FTD. Acta Neuropathol.

8. Benito E, Barco A (2015) The neuronal activity-driven transcriptome. Mol Neurobiol 51:1071–1088.

9. Berecki G, Helbig KL, Ware TL, Grinton B, Skraban CM, Marsh ED, Berkovic SF, Petrou S (2020) Novel Missense CACNA1G Mutations Associated with Infantile-Onset Developmental and Epileptic Encephalopathy. Int J Mol Sci 21.

10. Boulting GL, Durresi E, Ataman B, Sherman MA, Mei K, Harmin DA, Carter AC, Hochbaum DR, Granger AJ, Engreitz JM, Hrvatin S, Blanchard MR, Yang MG, Griffith EC, Greenberg ME (2021) Activity-dependent regulome of human GABAergic neurons reveals new patterns of gene regulation and neurological disease heritability. Nat Neurosci 24:437–448.

11. Burguete AS, Almeida S, Gao FB, Kalb R, Akins MR, Bonini NM (2015) GGGGCC microsatellite RNA is neuritically localized, induces branching defects, and perturbs transport granule function. Elife 4:e08881.

12. Burley S, Beccano-Kelly DA, Talbot K, Llana OC, Wade-Martins R (2022) Hyperexcitability in young iPSC-derived C9ORF72 mutant motor neurons is associated with increased intracellular calcium release. Scientific Reports 12.

13. Catanese A, Rajkumar S, Sommer D, Freisem D, Wirth A, Aly A, Massa-Lopez D, Olivieri A, Torelli F, Ioannidis V, Lipecka J, Guerrera IC, Zytnicki D, Ludolph A, Kabashi E, Mulaw MA, Roselli F, Bockers TM (2021) Synaptic disruption and CREB-regulated transcription are restored by K (+) channel blockers in ALS. EMBO Mol Med 13:e13131.

14. Cauchi RJ (2014) Gem depletion: amyotrophic lateral sclerosis and spinal muscular atrophy crossover. CNS Neurosci Ther 20:574–581.

15. Chen Y, Bang S, McMullen MF, Kazi H, Talbot K, Ho MX, Carlson G, Arnold SE, Ong WY, Kim SF (2017) Neuronal Activity-Induced Sterol Regulatory Element Binding Protein-1 (SREBP1) is Disrupted in Dysbindin-Null Mice-Potential Link to Cognitive Impairment in Schizophrenia. Mol Neurobiol 54:1699–1709.

16. Conesa A, Nueda MJ, Ferrer A, Talon M (2006) maSigPro: a method to identify significantly differential expression profiles in time-course microarray experiments. Bioinformatics 22:1096–1102.

17. Conlon EG, Lu L, Sharma A, Yamazaki T, Tang T, Shneider NA, Manley JL (2016) The C9ORF72 GGGGCC expansion forms RNA G-quadruplex inclusions and sequesters hnRNP H to disrupt splicing in ALS brains. Elife 5.

18. DeJesus-Hernandez M et al. (2011) Expanded GGGGCC hexanucleotide repeat in noncoding region of C9ORF72 causes chromosome 9p-linked FTD and ALS. Neuron 72:245–256.

19. Dell’Isola GB, Mencaroni E, Fattorusso A, Tascini G, Prontera P, Imperatore V, Di Cara G, Striano P, Verrotti A (2022) Expanding the genetic and clinical characteristics of Protocadherin 19 gene mutations. BMC Med Genomics 15:181.

20. Devlin AC, Burr K, Borooah S, Foster JD, Cleary EM, Geti I, Vallier L, Shaw CE, Chandran S, Miles GB (2015) Human iPSC-derived motoneurons harbouring TARDBP or C9ORF72 ALS mutations are dysfunctional despite maintaining viability. Nat Commun 6:5999.

21. Dobin A, Davis CA, Schlesinger F, Drenkow J, Zaleski C, Jha S, Batut P, Chaisson M, Gingeras TR (2013) STAR: ultrafast universal RNA-seq aligner. Bioinformatics 29:15–21.

22. Donnelly CJ, Zhang PW, Pham JT, Haeusler AR, Mistry NA, Vidensky S, Daley EL, Poth EM, Hoover B, Fines DM, Maragakis N, Tienari PJ, Petrucelli L, Traynor BJ, Wang J, Rigo F, Bennett CF, Blackshaw S, Sattler R, Rothstein JD (2013) RNA toxicity from the ALS/FTD C9ORF72 expansion is mitigated by antisense intervention. Neuron 80:415–428.

23. Ebbert MTW, Ross CA, Pregent LJ, Lank RJ, Zhang C, Katzman RB, Jansen-West K, Song Y, da Rocha EL, Palmucci C, Desaro P, Robertson AE, Caputo AM, Dickson DW, Boylan KB, Rademakers R, Ordog T, Li H, Belzil VV (2017) Conserved DNA methylation combined with differential frontal cortex and cerebellar expression distinguishes C9orf72-associated and sporadic ALS, and implicates SERPINA1 in disease. Acta Neuropathol 134:715–728.

24. Ebert DH, Greenberg ME (2013) Activity-dependent neuronal signalling and autism spectrum disorder. Nature 493:327–337.

25. Everett KV et al. (2007) Linkage and association analysis of CACNG3 in childhood absence epilepsy. Eur J Hum Genet 15:463–472.

26. Fernandopulle MS, Prestil R, Grunseich C, Wang C, Gan L, Ward ME (2018) Transcription Factor-Mediated Differentiation of Human iPSCs into Neurons. Curr Protoc Cell Biol 79:e51.

27. Flavell SW, Greenberg ME (2008) Signaling mechanisms linking neuronal activity to gene expression and plasticity of the nervous system. Annu Rev Neurosci 31:563–590.

28. Freibaum BD, Taylor JP (2017) The Role of Dipeptide Repeats in C9ORF72-Related ALS-FTD. Front Mol Neurosci 10:35.

29. Frick P, Sellier C, Mackenzie IRA, Cheng CY, Tahraoui-Bories J, Martinat C, Pasterkamp RJ, Prudlo J, Edbauer D, Oulad-Abdelghani M, Feederle R, Charlet-Berguerand N, Neumann M (2018) Novel antibodies reveal presynaptic localization of C9orf72 protein and reduced protein levels in C9orf72 mutation carriers. Acta Neuropathol Commun 6:72.

30. Fu J, Guo O, Zhen Z, Zhen J (2020) Essential Functions of the Transcription Factor Npas4 in Neural Circuit Development, Plasticity, and Diseases. Front Neurosci 14:603373.

31. Ghaffari LT, Trotti D, Haeusler AR (2023) Differential response of C9orf72 transcripts following neuronal depolarization. iScience 26.

32. Ghaffari LT, Trotti D, Haeusler AR, Jensen BK (2022) Breakdown of the central synapses in C9orf72-linked ALS/FTD. Front Mol Neurosci 15:1005112.

33. Ghiretti AE, Moore AR, Brenner RG, Chen LF, West AE, Lau NC, Van Hooser SD, Paradis S (2014) Rem2 is an activity-dependent negative regulator of dendritic complexity in vivo. J Neurosci 34:392–407.

34. Graw J (2009) Genetics of crystallins: cataract and beyond. Exp Eye Res 88:173–189.

35. Graw J (2017) From eyeless to neurological diseases. Exp Eye Res 156:5–9.

36. Guo JU, Ma DK, Mo H, Ball MP, Jang MH, Bonaguidi MA, Balazer JA, Eaves HL, Xie B, Ford E, Zhang K, Ming GL, Gao Y, Song H (2011) Neuronal activity modifies the DNA methylation landscape in the adult brain. Nat Neurosci 14:1345–1351.

37. Hadano S et al. (2001) A gene encoding a putative GTPase regulator is mutated in familial amyotrophic lateral sclerosis 2. Nat Genet 29:166–173.

38. Haeusler AR, Donnelly CJ, Periz G, Simko EA, Shaw PG, Kim MS, Maragakis NJ, Troncoso JC, Pandey A, Sattler R, Rothstein JD, Wang J (2014) C9orf72 nucleotide repeat structures initiate molecular cascades of disease. Nature 507:195–200.

39. Hahn A, Pensold D, Bayer C, Tittelmeier J, Gonzalez-Bermudez L, Marx-Blumel L, Linde J, Gross J, Salinas-Riester G, Lingner T, von Maltzahn J, Spehr M, Pieler T, Urbach A, Zimmer-Bensch G (2020) DNA Methyltransferase 1 (DNMT1) Function Is Implicated in the Age-Related Loss of Cortical Interneurons. Front Cell Dev Biol 8:639.

40. He Y, Casaccia-Bonnefil P (2008) The Yin and Yang of YY1 in the nervous system. J Neurochem 106:1493–1502.

41. Huang X et al. (2021) Human amyotrophic lateral sclerosis excitability phenotype screen: Target discovery and validation. Cell Rep 35:109224.

42. Hutnick LK, Golshani P, Namihira M, Xue Z, Matynia A, Yang XW, Silva AJ, Schweizer FE, Fan G (2009) DNA hypomethylation restricted to the murine forebrain induces cortical degeneration and impairs postnatal neuronal maturation. Hum Mol Genet 18:2875–2888.

43. Jackson JL, Finch NA, Baker MC, Kachergus JM, DeJesus-Hernandez M, Pereira K, Christopher E, Prudencio M, Heckman MG, Thompson EA, Dickson DW, Shah J, Oskarsson B, Petrucelli L, Rademakers R, van Blitterswijk M (2020) Elevated methylation levels, reduced expression levels, and frequent contractions in a clinical cohort of C9orf72 expansion carriers. Mol Neurodegener 15:7.

44. Jensen BK, Schuldi MH, McAvoy K, Russell KA, Boehringer A, Curran BM, Krishnamurthy K, Wen X, Westergard T, Ma L, Haeusler AR, Edbauer D, Pasinelli P, Trotti D (2020) Synaptic dysfunction induced by glycine-alanine dipeptides in C9orf72-ALS/FTD is rescued by SV2 replenishment. EMBO Mol Med 12:e10722.

45. Kato AS, Gill MB, Yu H, Nisenbaum ES, Bredt DS (2010) TARPs differentially decorate AMPA receptors to specify neuropharmacology. Trends Neurosci 33:241–248.

46. Kiernan MC (2009) Hyperexcitability, persistent Na+ conductances and neurodegeneration in amyotrophic lateral sclerosis. Exp Neurol 218:1–4.

47. Kiernan MC, Vucic S, Cheah BC, Turner MR, Eisen A, Hardiman O, Burrell JR, Zoing MC (2011) Amyotrophic lateral sclerosis. Lancet 377:942–955.

48. Kim S, Yu NK, Shim KW, Kim JI, Kim H, Han DH, Choi JE, Lee SW, Choi DI, Kim MW, Lee DS, Lee K, Galjart N, Lee YS, Lee JH, Kaang BK (2018) Remote Memory and Cortical Synaptic Plasticity Require Neuronal CCCTC-Binding Factor (CTCF). J Neurosci 38:5042–5052.

49. Kim TK, Hemberg M, Gray JM, Costa AM, Bear DM, Wu J, Harmin DA, Laptewicz M, Barbara-Haley K, Kuersten S, Markenscoff-Papadimitriou E, Kuhl D, Bito H, Worley PF, Kreiman G, Greenberg ME (2010) Widespread transcription at neuronal activity-regulated enhancers. Nature 465:182–187.

50. Laflamme C, McKeever PM, Kumar R, Schwartz J, Kolahdouzan M, Chen CX, You Z, Benaliouad F, Gileadi O, McBride HM, Durcan TM, Edwards AM, Healy LM, Robertson J, McPherson PS (2019) Implementation of an antibody characterization procedure and application to the major ALS/FTD disease gene C9ORF72. Elife 8.

51. Lall D, Lorenzini I, Mota TA, Bell S, Mahan TE, Ulrich JD, Davtyan H, Rexach JE, Muhammad A, Shelest O, Landeros J, Vazquez M, Kim J, Ghaffari L, O’Rourke JG, Geschwind DH, Blurton-Jones M, Holtzman DM, Sattler R, Baloh RH (2021) C9orf72 deficiency promotes microglial-mediated synaptic loss in aging and amyloid accumulation. Neuron 109:2275–2291 e2278.

52. Laszlo ZI, Hindley N, Sanchez Avila A, Kline RA, Eaton SL, Lamont DJ, Smith C, Spires-Jones TL, Wishart TM, Henstridge CM (2022) Synaptic proteomics reveal distinct molecular signatures of cognitive change and C9ORF72 repeat expansion in the human ALS cortex. Acta Neuropathol Commun 10:156.

53. Lee YB, Chen HJ, Peres JN, Gomez-Deza J, Attig J, Stalekar M, Troakes C, Nishimura AL, Scotter EL, Vance C, Adachi Y, Sardone V, Miller JW, Smith BN, Gallo JM, Ule J, Hirth F, Rogelj B, Houart C, Shaw CE (2013) Hexanucleotide repeats in ALS/FTD form length-dependent RNA foci, sequester RNA binding proteins, and are neurotoxic. Cell Rep 5:1178–1186.

54. Liang H, Ward WF, Jang YC, Bhattacharya A, Bokov AF, Li Y, Jernigan A, Richardson A, Van Remmen H (2011) PGC-1alpha protects neurons and alters disease progression in an amyotrophic lateral sclerosis mouse model. Muscle Nerve 44:947–956.

55. Liao Y, Smyth GK, Shi W (2019) The R package Rsubread is easier, faster, cheaper and better for alignment and quantification of RNA sequencing reads. Nucleic Acids Res 47:e47.

56. Ling SC, Polymenidou M, Cleveland DW (2013) Converging mechanisms in ALS and FTD: disrupted RNA and protein homeostasis. Neuron 79:416–438.

57. Liu H, Bell K, Herrmann A, Arnhold S, Mercieca K, Anders F, Nagel-Wolfrum K, Thanos S, Prokosch V (2022) Crystallins Play a Crucial Role in Glaucoma and Promote Neuronal Cell Survival in an In Vitro Model Through Modulating Muller Cell Secretion. Invest Ophthalmol Vis Sci 63:3.

58. Love MI, Huber W, Anders S (2014) Moderated estimation of fold change and dispersion for RNA-seq data with DESeq2. Genome Biol 15:550.

59. Mackenzie IR, Arzberger T, Kremmer E, Troost D, Lorenzl S, Mori K, Weng SM, Haass C, Kretzschmar HA, Edbauer D, Neumann M (2013) Dipeptide repeat protein pathology in C9ORF72 mutation cases: clinico-pathological correlations. Acta Neuropathol 126:859–879.

60. Mao JJ, Katayama S, Watanabe C, Harada Y, Noda K, Yamamura Y, Nakamura S (2001) The relationship between alphaB-crystallin and neurofibrillary tangles in Alzheimer’s disease. Neuropathol Appl Neurobiol 27:180–188.

61. Mao YW, Liu JP, Xiang H, Li DW (2004) Human alphaA- and alphaB-crystallins bind to Bax and Bcl-X (S) to sequester their translocation during staurosporine-induced apoptosis. Cell Death Differ 11:512–526.

62. Menon P, Higashihara M, van den Bos M, Geevasinga N, Kiernan MC, Vucic S (2020) Cortical hyperexcitability evolves with disease progression in ALS. Ann Clin Transl Neurol 7:733–741.

63. Moore AR, Richards SE, Kenny K, Royer L, Chan U, Flavahan K, Van Hooser SD, Paradis S (2018) Rem2 stabilizes intrinsic excitability and spontaneous firing in visual circuits. Elife 7.

64. Mori K, Lammich S, Mackenzie IR, Forne I, Zilow S, Kretzschmar H, Edbauer D, Janssens J, Kleinberger G, Cruts M, Herms J, Neumann M, Van Broeckhoven C, Arzberger T, Haass C (2013) hnRNP A3 binds to GGGGCC repeats and is a constituent of p62-positive/TDP43-negative inclusions in the hippocampus of patients with C9orf72 mutations. Acta Neuropathol 125:413–423.

65. Nueda MJ, Tarazona S, Conesa A (2014) Next maSigPro: updating maSigPro bioconductor package for RNA-seq time series. Bioinformatics 30:2598–2602.

66. Opsomer R, Contino S, Perrin F, Gualdani R, Tasiaux B, Doyen P, Vergouts M, Vrancx C, Doshina A, Pierrot N, Octave JN, Gailly P, Stanga S, Kienlen-Campard P (2020) Amyloid Precursor Protein (APP) Controls the Expression of the Transcriptional Activator Neuronal PAS Domain Protein 4 (NPAS4) and Synaptic GABA Release. eNeuro 7.

67. Osenberg S, Karten A, Sun J, Li J, Charkowick S, Felice CA, Kritzer M, Nguyen MVC, Yu P, Ballas N (2018) Activity-dependent aberrations in gene expression and alternative splicing in a mouse model of Rett syndrome. Proc Natl Acad Sci U S A 115:E5363–E5372.

68. Pabian-Jewula S, Bragiel-Pieczonka A, Rylski M (2022) Ying Yang 1 engagement in brain pathology. J Neurochem 161:236–253.

69. Pancho A, Aerts T, Mitsogiannis MD, Seuntjens E (2020) Protocadherins at the Crossroad of Signaling Pathways. Front Mol Neurosci 13:117.

70. Pollina EA, Gilliam DT, Landau AT, Lin C, Pajarillo N, Davis CP, Harmin DA, Yap EL, Vogel IR, Griffith EC, Nagy MA, Ling E, Duffy EE, Sabatini BL, Weitz CJ, Greenberg ME (2023) A NPAS4-NuA4 complex couples synaptic activity to DNA repair. Nature 614:732–741.

71. Renton AE et al. (2011) A hexanucleotide repeat expansion in C9ORF72 is the cause of chromosome 9p21-linked ALS-FTD. Neuron 72:257–268.

72. Rienecker KDA, Poston RG, Saha RN (2020) Merits and Limitations of Studying Neuronal Depolarization-Dependent Processes Using Elevated External Potassium. ASN Neuro 12:1759091420974807.

73. Rosenblum LT, Trotti D (2017) EAAT2 and the Molecular Signature of Amyotrophic Lateral Sclerosis. Adv Neurobiol 16:117–136.

74. Roussos P, Guennewig B, Kaczorowski DC, Barry G, Brennand KJ (2016) Activity-Dependent Changes in Gene Expression in Schizophrenia Human-Induced Pluripotent Stem Cell Neurons. JAMA Psychiatry 73:1180–1188.

75. Rudenko A, Dawlaty MM, Seo J, Cheng AW, Meng J, Le T, Faull KF, Jaenisch R, Tsai LH (2013) Tet1 is critical for neuronal activity-regulated gene expression and memory extinction. Neuron 79:1109–1122.

76. Sareen D, O’Rourke JG, Meera P, Muhammad AK, Grant S, Simpkinson M, Bell S, Carmona S, Ornelas L, Sahabian A, Gendron T, Petrucelli L, Baughn M, Ravits J, Harms MB, Rigo F, Bennett CF, Otis TS, Svendsen CN, Baloh RH (2013) Targeting RNA foci in iPSC-derived motor neurons from ALS patients with a C9ORF72 repeat expansion. Sci Transl Med 5:208ra149.

77. Satoh J, Yamamoto Y, Kitano S, Takitani M, Asahina N, Kino Y (2014) Molecular network analysis suggests a logical hypothesis for the pathological role of c9orf72 in amyotrophic lateral sclerosis/frontotemporal dementia. J Cent Nerv Syst Dis 6:69–78.

78. Schanz O, Bageac D, Braun L, Traynor BJ, Lehky TJ, Floeter MK (2016) Cortical hyperexcitability in patients with C9ORF72 mutations: Relationship to phenotype. Muscle Nerve 54:264–269.

79. Shi Y et al. (2018) Haploinsufficiency leads to neurodegeneration in C9ORF72 ALS/FTD human induced motor neurons. Nat Med 24:313–325.

80. Shibuya K, Otani R, Suzuki YI, Kuwabara S, Kiernan MC (2022) Neuronal Hyperexcitability and Free Radical Toxicity in Amyotrophic Lateral Sclerosis: Established and Future Targets. Pharmaceuticals (Basel) 15.

81. Su Y, Shin J, Zhong C, Wang S, Roychowdhury P, Lim J, Kim D, Ming GL, Song H (2017) Neuronal activity modifies the chromatin accessibility landscape in the adult brain. Nat Neurosci 20:476–483.

82. Sullivan PM, Zhou X, Robins AM, Paushter DH, Kim D, Smolka MB, Hu F (2016) The ALS/FTLD associated protein C9orf72 associates with SMCR8 and WDR41 to regulate the autophagy-lysosome pathway. Acta Neuropathol Commun 4:51.

83. Thau N, Knippenberg S, Korner S, Rath KJ, Dengler R, Petri S (2012) Decreased mRNA expression of PGC-1alpha and PGC-1alpha-regulated factors in the SOD1G93A ALS mouse model and in human sporadic ALS. J Neuropathol Exp Neurol 71:1064–1074.

84. Varghese M, Zhao W, Trageser KJ, Pasinetti GM (2020) Peroxisome Proliferator Activator Receptor Gamma Coactivator-1alpha Overexpression in Amyotrophic Lateral Sclerosis: A Tale of Two Transgenics. Biomolecules 10.

85. Verheul TCJ, van Hijfte L, Perenthaler E, Barakat TS (2020) The Why of YY1: Mechanisms of Transcriptional Regulation by Yin Yang 1. Front Cell Dev Biol 8:592164.

86. Vucic S, Nicholson GA, Kiernan MC (2008) Cortical hyperexcitability may precede the onset of familial amyotrophic lateral sclerosis. Brain 131:1540–1550.

87. Vucic S, Ziemann U, Eisen A, Hallett M, Kiernan MC (2013) Transcranial magnetic stimulation and amyotrophic lateral sclerosis: pathophysiological insights. J Neurol Neurosurg Psychiatry 84:1161–1170.

88. Vucic S, Pavey N, Haidar M, Turner BJ, Kiernan MC (2021) Cortical hyperexcitability: Diagnostic and pathogenic biomarker of ALS. Neurosci Lett 759:136039.

89. Wainger BJ, Cudkowicz ME (2015) Cortical Hyperexcitability in Amyotrophic Lateral Sclerosis: C9orf72 Repeats. JAMA Neurol 72:1235–1236.

90. Weskamp K, Tank EM, Miguez R, McBride JP, Gomez NB, White M, Lin Z, Gonzalez CM, Serio A, Sreedharan J, Barmada SJ (2020) Shortened TDP43 isoforms upregulated by neuronal hyperactivity drive TDP43 pathology in ALS. J Clin Invest 130:1139–1155.

91. Westergard T, McAvoy K, Russell K, Wen X, Pang Y, Morris B, Pasinelli P, Trotti D, Haeusler A (2019) Repeat-associated non-AUG translation in C9orf72-ALS/FTD is driven by neuronal excitation and stress. EMBO Mol Med 11.

92. Xiao S, McKeever PM, Lau A, Robertson J (2019) Synaptic localization of C9orf72 regulates post-synaptic glutamate receptor 1 levels. Acta Neuropathol Commun 7:161.

93. Yap EL, Greenberg ME (2018) Activity-Regulated Transcription: Bridging the Gap between Neural Activity and Behavior. Neuron 100:330–348.

94. Zhang M, Tartaglia MC, Moreno D, Sato C, McKeever P, Weichert A, Keith J, Robertson J, Zinman L, Rogaeva E (2017) DNA methylation age-acceleration is associated with disease duration and age at onset in C9orf72 patients. Acta Neuropathol 134:271–279.

95. Zhang SJ, Buchthal B, Lau D, Hayer S, Dick O, Schwaninger M, Veltkamp R, Zou M, Weiss U, Bading H (2011) A signaling cascade of nuclear calcium-CREB-ATF3 activated by synaptic NMDA receptors defines a gene repression module that protects against extrasynaptic NMDA receptor-induced neuronal cell death and ischemic brain damage. J Neurosci 31:4978–4990.

96. Zhao W, Varghese M, Yemul S, Pan Y, Cheng A, Marano P, Hassan S, Vempati P, Chen F, Qian X, Pasinetti GM (2011) Peroxisome proliferator activator receptor gamma coactivator-1alpha (PGC-1alpha) improves motor performance and survival in a mouse model of amyotrophic lateral sclerosis. Mol Neurodegener 6:51.

